# From HeLa to human blood: extending single-cell proteomics to the smallest immune cells

**DOI:** 10.1101/2025.11.10.687556

**Authors:** Ivo A. Hendriks, Sara C. Buch-Larsen, Martin Rykær, Maico Y. Lechner, Anders H. Kverneland, Tabiwang N. Arrey, Daniel Hermanson, Eugen Damoc, Jesper V. Olsen

## Abstract

Single-cell proteomics (SCP) holds the promise of decoding cellular heterogeneity at the functional level, yet achieving deep and reproducible proteome coverage from individual cells has remained a formidable challenge. Here, we established a high-sensitivity, label-free SCP platform that surpasses previous limits in depth and reproducibility. By integrating optimized low-input sample processing, refined liquid chromatography, and the Orbitrap Astral Zoom mass spectrometer, our approach routinely quantified over 7,000 proteins per individual HeLa cell, capturing thousands of low-abundance proteins that eluded prior SCP studies. Applied to very small human peripheral blood mononuclear cells (PBMCs), we identified up to 4,000 proteins per cell, including key markers distinguishing monocytes, T cells, and activated lymphocytes within heterogeneous populations, underscoring that single-cell proteomics can now directly elucidate clinically relevant primary samples with both depth and precision.

## INTRODUCTION

The recent technological advances made in single-cell proteomics (SCP) have revolutionized our ability to interrogate the proteome at the resolution of individual cells, offering unprecedented insights into cellular heterogeneity, functional diversity, and dynamics of biological processes that are often masked in bulk analyses. Unlike traditional bulk proteomics, which averages protein abundances across large populations of cells, SCP enables the direct quantification and characterization of proteins, including their post-translational modifications, within single cells. This granular approach is critical for understanding complex biological phenomena such as cellular differentiation, disease progression, immune responses, and drug resistance, as well as for identifying rare cell types that may play pivotal roles in health and disease.

The last few years have witnessed remarkable progress in both the sensitivity and throughput of SCP, driven by innovations in sample preparation workflows and mass spectrometry (MS) acquisition strategies (Lin et al., 2025). The primary challenge in SCP lies in the minute amount of protein available from individual cells, necessitating highly efficient and low-loss sample processing workflows coupled with ultra-sensitive MS detection. The SCP field has seen a surge in the development of miniaturized and automated sample preparation platforms. This includes the nanoPOTS (Nanodroplet Processing in One pot for Trace Samples) and the more recent nested NanoPOTS (N2) chip, which leverage microfluidic and nanodroplet technologies to minimize sample loss, reduce reaction volumes, and enable the parallel processing of hundreds of single cells (Woo et al., 2021). These platforms have demonstrated robust quantification of thousands of proteins per cell, with the latest iterations, such as the proteoCHIP EVO 96 and the Chip-Tip workflow, pushing the boundaries further by enabling the identification of more than 5,000 proteins from individual cells in a high-throughput and nearly loss-less manner (Ye et al., 2025). Parallel to advances in sample preparation, highly sensitive and fast sequencing mass spectrometers such as the Orbitrap Astral and the timsTOF mass spectrometers have been introduced to meet the demands of SCP. Data-independent acquisition (DIA) has emerged as a preferred MS acquisition strategy due to its reproducibility, comprehensive coverage, and suitability for low-input samples. Optimized narrow-window DIA workflows (Guzman et al., 2024) often coupled with short liquid chromatography (LC) nanoflow gradients and spectral library-free data analysis, have enabled the identification of thousands of proteins from single cells with high reproducibility and quantitative accuracy. Additionally, isobaric labeling techniques, such as those employed in SCoPE-MS (Budnik et al., 2018), utilizing tandem mass tags (TMT) and carrier channels to boost signal intensity and facilitate multiplexed analysis, further enhancing throughput and sensitivity (Ye et al., 2022).

Despite these advances, SCP still faces challenges related to sample loss, data analysis, and the need for further improvements in sensitivity and throughput. Ongoing efforts in miniaturization, automation, and the development of robust computational tools are essential for overcoming these limitations and realizing the full potential of SCP in both basic and translational research. Using a pulsed-SILAC strategy, we have recently demonstrated that single-cell proteomics can be used to study the relationship between the turnover and concentrations of certain proteins compared to cell size in human cells (Sabatier et al., 2025).

Protein content in most cell types across the tree of life is in the ranges of 15-35% of the total cell volume with cellular protein concentration typically around 200-300 mg/mL. This cellular constraint is thought to be a result of evolutionary pressure to optimize metabolic efficiency (Brown, 1991). A single cell’s total protein mass can therefore vary significantly with cell size, with larger cells having more protein copies. However, not all proteins scale equally; while some like histones are diluted, others are concentrated as cells grow. Cell size is a major source of proteome heterogeneity, and consequently cell size effects have a confounding role in SCP. Even though this effect is obfuscated in the majority of published SCP datasets since they consist of relatively homogenous cell types (e.g. epithelial cells like HeLa) with little variation in cell sizes, proteome coverage in single-cell proteomics generally scales with the size of the cells studied (Bubis et al., 2025) with significantly lower number of proteins identified and quantified in smaller cells like blood cells (Furtwangler et al., 2025). This is likely due to technical factors such as raw MS sensitivity, as smaller cells contain fewer copies of each protein, making it harder to detect low-abundance proteins from smaller cells. Moreover, even minor losses during single cell proteomics sample preparation disproportionately affect smaller cells. Therefore, further improvements in sample preparation and MS sensitivity is required to study smaller cell types such as immune cells by single-cell proteomics. Here, we provide a systematic optimization of a label-free SCP workflow for high-sensitivity single-cell proteomics, with a particular emphasis on improving single-cell proteome coverage by leveraging improved sample preparation strategies and enhanced mass spectrometry acquisition methods on the Thermo Scientific^TM^ Orbitrap™ Astral™ Zoom mass spectrometer. Based on these technological innovations, we demonstrate routine coverage of >7,000 proteins in single HeLa cells, and ∼4,000 proteins from individual small cells prepared from human peripheral blood mononuclear cells (PBMCs), providing proteome coverage of >50% of the adaptive immunity pathways. This SCP technology outlines future directions for advancing single-cell proteomics towards routine, large-scale applications in biology and medicine.

## RESULTS

### Systematic optimization of LC-MS conditions for low load

Mass spectrometry (MS)-based single-cell proteomics (SCP) is an emerging field, with a plethora of sample preparation and technological improvements resulting in enhanced depth of sequencing from highly limited starting material. Here, we endeavored to systematically assess, optimize, and apply cutting-edge methodology and technology in order to maximize insight that can be gleaned from analysis of the smallest single cells. To this end, we first set out to optimize liquid chromatography (LC) and MS methodology for analysis of low amounts of peptides derived from HeLa digests (Fig. 1A), using a Thermo Scientific^TM^ Vanquish™ Neo UHPLC System (Vanquish) and narrow-window data-independent acquisition (nDIA) on a Thermo Scientific™ Orbitrap™ Astral™ MS (Astral).

**Figure 1.**
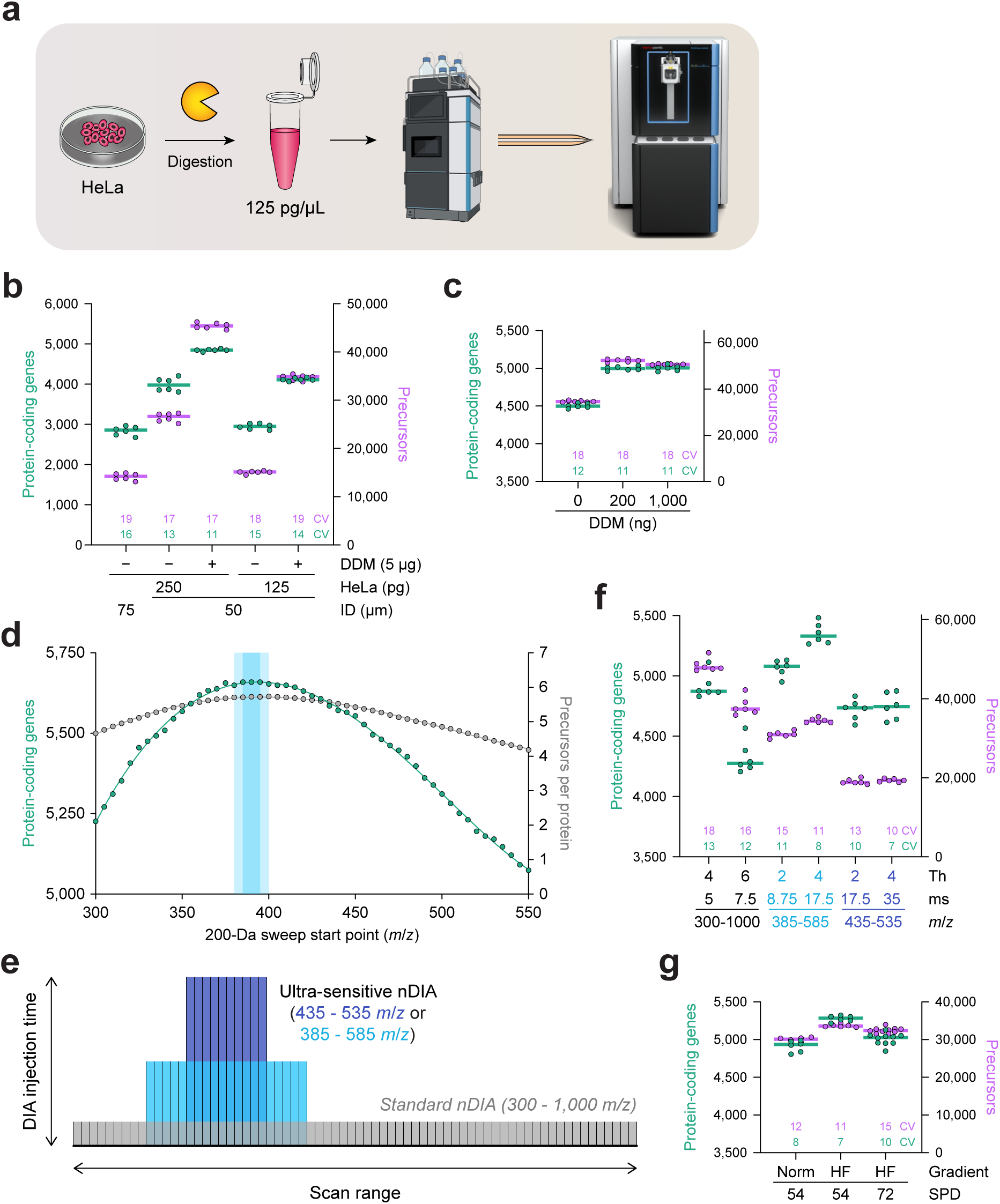
Optimization of low-input LC-MS methodology for enhanced protein identification rates. (A) Schematic overview of the experimental design. HeLa cells were lysed, digested with Lys-C and trypsin, and purified HeLa peptides were diluted to a concentration of 125 pg/µL prior to analysis using a Vanquish HPLC, laser-pulled and in-house packed analytical columns, and an Astral mass spectrometer. (B) Number of protein-coding genes (green, left axis) and precursors (purple, right axis) identified, when comparing different column inner diameter, HeLa load, and the addition of n-dodecyl β-D-maltoside (DDM). Bars represent median, the numbers above the x-axis represent median coefficient of variation percentage, *n* = 6 technical replicates. “ID”; internal column diameter. Data analysis using DIA-NN v1.9.2 without matching-between-runs. (C) As **B**, but comparing different amounts of DDM injected for analysis of 250 pg HeLa. (D) Number of protein-coding genes (green, left axis) and precursors per protein (grey, right axis) identified when restricting identification to a 200 Da window, e.g. from 300 – 500 *m*/*z* when starting at 300 *m*/*z*, based on the data from **B** that was recorded from 300 – 1,000 *m*/*z*. Blue highlighting indicates the local optimum range for identification of the most protein-coding genes. (E) Visualization of the ultra-sensitive narrow-window DIA (nDIA) concept, where sampling is restricted to a narrower but more proteotypic *m/z* range, while increasing maximum DIA ion injection time to enhance sensitivity. (F) As **B**, but comparing different ultra-sensitive nDIA methods for analysis of 250 pg HeLa. “ms”; maximum DIA ion injection time in milliseconds. (G) As **B**, but evaluating different gradient lengths and throughput speeds for analysis of 250 pg HeLa. *n* = 6-10 technical replicates. “Norm”; standard steady-flow chromatography, “HF”; method employing Higher Flow at the start and end of the gradient to maximize effective peptide elution time, “SPD”; samples per day.

To optimize LC-MS settings for SCP, we first analyzed dilutions of a bulk tryptic digest of HeLa lysate. We compared analysis of 250 or 125 pg peptide digest (peptide content roughly equivalent to one or half of a single HeLa cell), in the absence or presence of the MS-compatible non-ionic detergent n-dodecyl β-D-maltoside (DDM), and using laser-pulled in-house packed C18 reversed-phase columns with an internal diameter (ID) of either 50 or the standard 75 µm (Fig. 1B and S1A). The different column diameters were tested to take advantage of the fact that the electrospray process is concentration dependent, and since the ideal flow rate for nanoLC is proportional to the column diameter, narrower columns support lower flow rates resulting in greater sensitivity (Zheng et al., 2023). Notably, LC-MS analysis was performed at comparable backpressure, with flow rate at 125 or 250 nL/min for the 50 and 75 µm columns, respectively. The number of identified precursors almost doubled when using a 50 µm ID column, and was further increased by the addition of DDM to the runs. Similar increases in identifications were observed for protein-coding genes (referred to as proteins hereafter), with a 50 µm ID column facilitating identification of as many proteins from 125 pg, as a 75 µm ID column with 250 pg (Fig. 1B). Along the same lines, and on the same 50 µm ID column, addition of DDM resulted in as many proteins profiled from 125 pg as compared to 250 pg without DDM. Sample loss due to adsorption at hydrophobic surfaces is a major bottleneck in SCP, but DDM can mitigate this by improving recovery of hydrophobic peptides, and thereby enhance signal intensity in MS-based SCP (Tsai et al., 2021). We reduced the amount of DDM by up to 25-fold (Fig. 1C and S1B), and found that even the smallest amount tested here was sufficient to limit peptide loss and enhance identification rates, overall allowing identification of ∼5,000 proteins and ∼50,000 precursors.

While we employed a conventional broad-range DIA-MS/MS scanning method (300-1000 m/z) at first, we reasoned that restricting the scan range while increasing the allowed injection time per window could enhance sensitivity while retaining proteotypicity. We scrutinized our first data to extract 200 m/z DIA-MS/MS scan ranges and determined the number of proteins within each of the scan ranges (Fig. 1D). We found that starting the scan range near ∼385 m/z resulted in maximum number of proteins and precursors-per-protein. To validate this, we set up several ultra-sensitive narrow-window DIA (nDIA) methods (Fig. 1E), in all cases keeping the overall scan cycle time constant at ∼1 second, and evaluated their performance on 250 pg HeLa (Fig. 1F and S1C). Indeed, although we observed a drop in the number of precursors identified when using a narrow scan range (385-585 m/z with 4-Th nDIA-MS/MS isolation windows and allowing 17.5 ms injection times), the number of proteins increased from ∼4,900 to ∼5,300, while also resulting in considerably lower coefficient of variation (Fig. 1F). Reducing the DIA-MS/MS scan range to only 100 m/z was not as efficient, and for the 200 m/z method we found that 4 Th windows were superior to 2 Th windows.

The number of samples per day (SPD) that can be analyzed is a pivotal metric to high-throughput and especially single-cell proteomics, as numerous single cells need to be measured to achieve sufficient statistical power. We designed a chromatographic method for the Vanquish Neo LC whereby we initiated the gradient at high flow and pressure, and then gradually reduced the flow prior to elution of peptides, effectively reducing column delay from 6 to 2 min. This allowed us to acquire essentially the same number of precursors and proteins in a 15 min gradient (72 SPD) compared to a 20 min gradient (54 SPD), with only a slightly increased coefficient of variation (Fig. 1G and S1D).

Several other MS-technical parameters were evaluated, including scanning loop control and fragment ranges (Supplementary Note 1). Together, these optimizations in LC-MS methodology – encompassing column selection, surfactant-assisted peptide recovery, refined scan ranges, and accelerated chromatographic gradients – substantially improved proteome coverage and throughput from minimal sample inputs, thereby establishing a robust foundation for subsequent experiments and single-cell proteomic analyses.

### Evaluating the Orbitrap Astral Zoom MS for enhanced sensitivity and proteome depth

Recently, the Orbitrap Astral Zoom MS (Astral Zoom) was introduced commercially. This instrument boasts several new features including pre-accumulation (Harking et al., 2025), higher acquisition rate of up to 270 Hz, and enhanced spectral processing (Guzman et al., 2025). Further, the Astral Zoom has a low input mode (LIM), which increases sensitivity for low input samples. For this study, we had access to a prototype Astral Zoom that allowed us to toggle these features individually, and thus perform a fair comparison between Astral and Astral Zoom (Fig. 2A).

**Figure 2.**
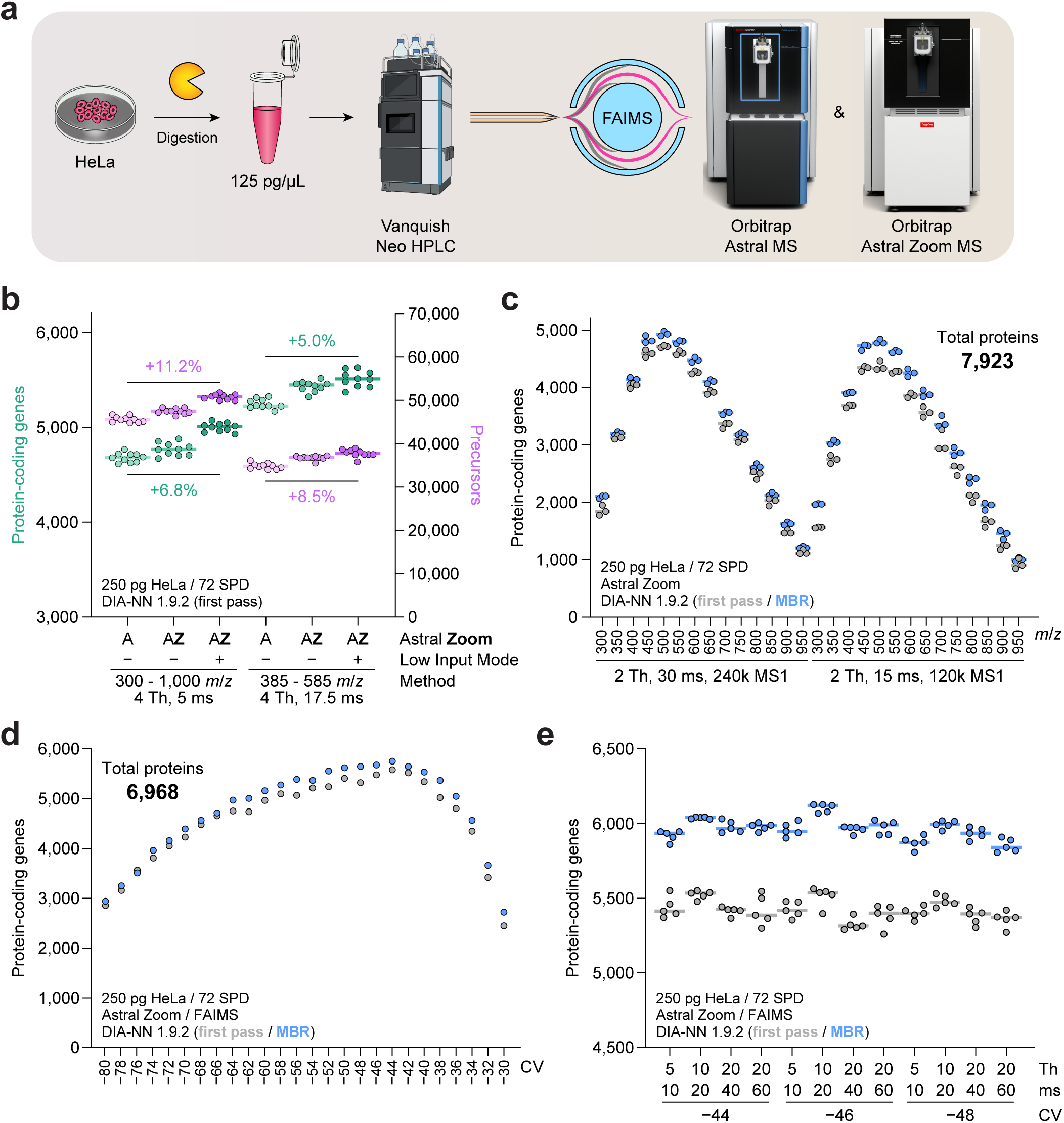
Evaluation of the Orbitrap Astral Zoom MS for low input analysis. (A) Schematic overview of the experimental design. HeLa cells were lysed, digested with Lys-C and trypsin, and purified HeLa peptides were diluted to a concentration of 125 pg/µL prior to analysis using a Vanquish HPLC, laser-pulled and in-house packed analytical columns, and an Orbitrap Astral Zoom prototype mass spectrometer. (B) Number of protein-coding genes (red, left axis) and precursors (blue, right axis) identified, when comparing “Astral” mode with “Astral Zoom” mode, with the latter either in regular or Low Input mode. Bars represent median, percentage increases of median values are indicated, *n* = 10 technical replicates. Data analysis using DIA-NN v1.9.2 without matching-between-runs. (C) Gas-phase fractionation experiment, wherein precursors are isolated from a 50 *m/z* range starting at the *m/z* indicated on the x-axis. Gray and blue dots correspond to the number of identified protein-coding genes with data analysis using DIA-NN in first pass mode and MBR mode, respectively. *n* = 3 technical replicates, total number of protein-coding genes identified cumulatively across all runs is indicated. (D) As **C**, but evaluating a range of FAIMS compensation voltages (CV), using a single constant CV across the entire gradient. *N* = 1 technical replicate. (E) As **D**, but investigating different combinations of DIA window size in Thomson (“Th”) and DIA ion injection time in milliseconds (“ms”) in combination with three different CVs. *n* = 5 technical replicates.

First, we evaluated performance when analyzing 250 pg HeLa, for both ultra-sensitive nDIA and standard methods (Fig. 2B and S2). At the precursor level, we observed an increase of 8.5% and 11.2% when running the Astral Zoom with LIM, and similarly the number of identified proteins increased by 5.0% and 6.8%. Coefficient of variation was similar or slightly lower with the Astral Zoom (Fig. S2), and significantly lower when using the ultra-sensitive nDIA method. We performed a more in-depth evaluation of the LIM, which is detailed in Supplementary Note 2.

We investigated optimal scan ranges for ultra-sensitive nDIA, initially approximating these empirically by determining the number of proteins within 200 m/z DIA-MS/MS scan ranges (Fig. 1D). To further validate this, we performed a gas-phase fractionation (GPF) experiment, where the scan range of each run was limited to 50 m/z, using two different nDIA acquisition speeds (Fig. 2C and S3). Analyzing 250 pg HeLa per LC-MS run, we found that the 450-500 m/z and 500-550 m/z fractions resulted in the highest identification rates, with ∼5,000 proteins identified, and consistent with an optimal scan range centered between 475 and 525 m/z. Moreover, cumulatively ∼110,000 unique peptides and ∼7,900 proteins could be identified across the entire GPF experiment (Fig. S3A-C), without ever injecting more than 250 pg per run.

An alternative strategy of fractionation in the gas-phase is through application of high-field asymmetric ion mobility spectrometry (FAIMS) using different compensation voltages (CVs), which impart selectivity mediated by mass-to-charge and mobility properties of precursors. We performed a FAIMS CV sweep by analyzing 250 pg of HeLa using single CVs in steps of 2 V ranging from −80 to −30 V (Fig. 2D and S4), both to determine an optimal CV range, and to investigate complementarity of a multi-CV fractionation approach. Although there was some variance between data analysis with DIA-NN and Spectronaut, as well as at the precursor and protein levels, CVs of −44, −46, and −48 V were proximate to the optimum (Fig. 2D and S4A-C). Cumulatively, we identified ∼90,000 unique peptides and ∼7,900 proteins using Spectronaut (Fig. S4B-C). Interestingly, there was a direct linear correlation between the number of observed tryptic missed cleavages in identified peptides and the compensation voltage applied (Fig. S4D). In terms of CV fraction complementarity, neighboring CVs demonstrated a high Pearson correlation (Fig. S4E), suggesting that large jumps in CV are necessary for maximum orthogonality. Finally, to determine an optimal CV to use for further experiments, we performed replicate analyses using CVs of −44, −46, and −48 V and four different combinations of isolation window and maximum injection times (Fig. 2E and S5). Based on this, we did not observe large differences between the CVs, but found that −46 V with a 10 Th / 20 ms method yielded the highest number of identifications. However, coefficient of variation was the lowest using a 20 Th / 40 ms method (Fig. S5B-C).

Collectively, these experiments demonstrate that the Orbitrap Astral Zoom MS, particularly when operated in low input mode, provides measurable gains in proteome coverage, sensitivity, and reproducibility, thereby advancing the analytical capabilities of low-input proteomics.

### Defining the limit of detection and quantitative precision at ultra-low input

The amount of protein derived from a single HeLa cell is typically assumed to be in the range of 200 to 250 pg (Dolgalev et al., 2023; Kulak et al., 2014). However, even within an asynchronous population of HeLa cells, there are considerable differences in cell size and shape. Moreover, many other cell types can have dramatically different sizes. Total protein amount per cell increases proportionally with cell size, and protein coverage in SCP scales positively with cell size due to both more total protein and higher abundance of detectable proteins, in addition to technical factors such as improved extraction and detection sensitivity. Thus, to anticipate analysis of smaller and more challenging cells, we performed a series of HeLa dilution experiments to determine the limit of detection of our LC-MS setup, as well as to further optimize our methodology in the context of ultra-low input amounts.

Utilizing the low-input methodology we established for analysis without and with FAIMS (Figs. 1 and 2), we compared performance of the methods over a dilution series starting at 100 pg and going down to as little as 0.5 pg (Fig. 3A-B and S6A-B). Down to 50 pg, we did not observe a notable difference between the methods when performing a first pass analysis using DIA-NN (Fig. 3A), demonstrating that both no-FAIMS and FAIMS approaches can be efficiently applied to loads above 50 pg. Of note, the FAIMS approaches performed slightly better when analyzing the data using Spectronaut (Fig. 3B), indicating that different software can lead to distinct conclusions. Consistently, when analyzing 25 pg or less, the FAIMS methodology started to outperform the no-FAIMS methodology, with up to ∼2-fold increases in the number of identified proteins at 0.5-2.5 pg loads. Quantitatively, coefficients of variation were highly comparable between the ultra-sensitive nDIA method and the FAIMS-specific MS acquisition method using 20 Th DIA windows and maximum allowed 40 ms injection time. To illustrate the effect of FAIMS in the context of ultra-low loads, we investigated the charge state distribution within the total ion current in two representative 1 pg HeLa runs, either with no-FAIMS analysis or FAIMS analysis (Fig. 3C). Without FAIMS, the vast majority of ions were singly charged and likely chemical background noise, with a high polysiloxane presence (Olsen et al., 2004). With FAIMS operated at optimal single CV of −46 V, singly charged precursors were almost entirely abolished, leaving predominantly doubly and triply charged peptide precursors.

**Figure 3.**
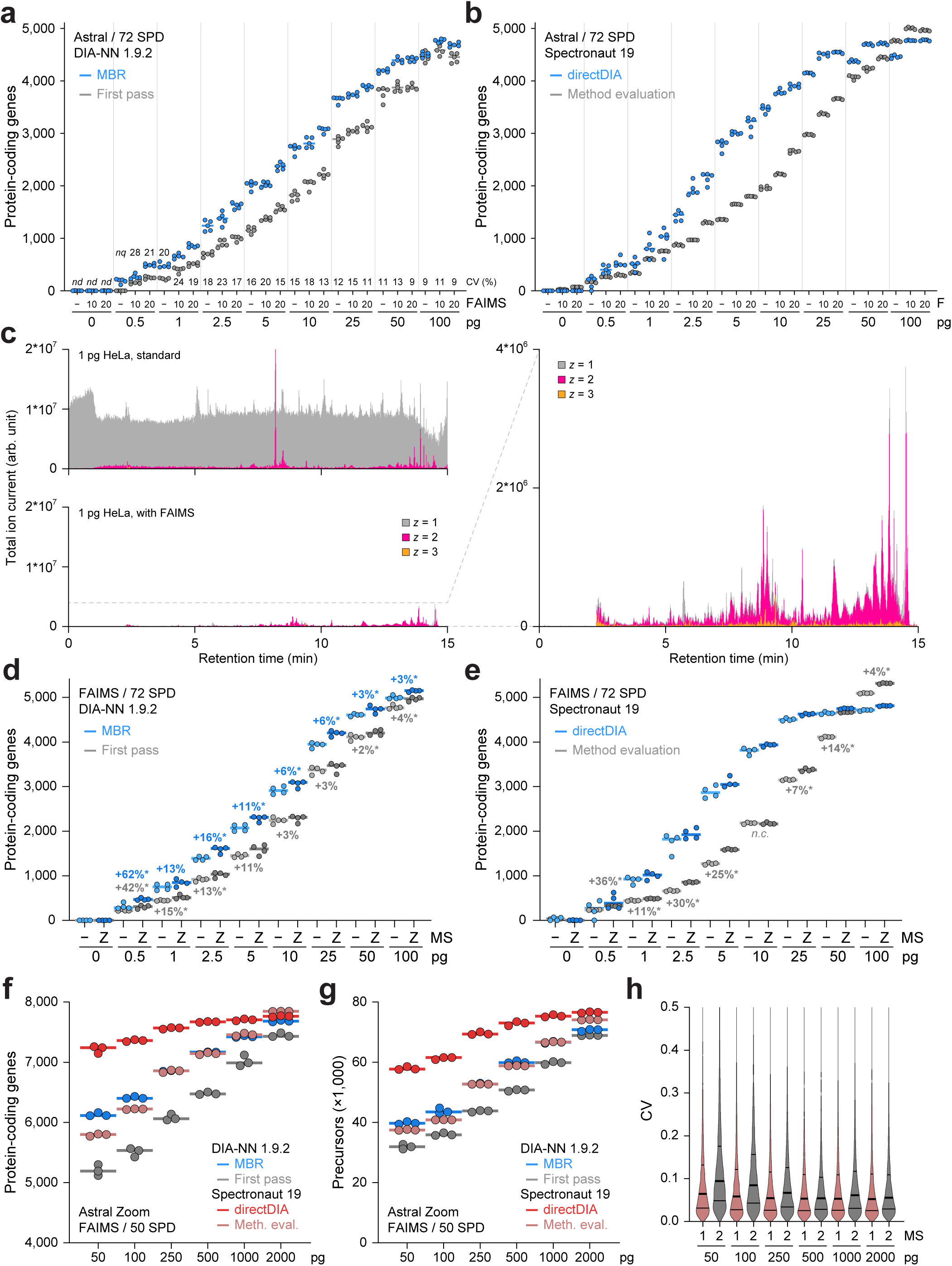
Orbitrap Astral Zoom MS and FAIMS are superior for ultra-low load analysis. (A) Dilution series experiment, ranging from 0.5 pg to 100 pg of injected HeLa lysate, comparing standard analysis to FAIMS analysis. All injections were performed in ascending amounts, starting with a blank to ensure negligible sample carry-over. Number of identified protein-coding genes are indicated, bars represent median, with gray and blue corresponding to data analysis using DIA-NN v1.9.2 in first pass mode and MBR mode, respectively. Numbers above the x-axis represent median coefficient of variation percentage, *n* = 5 technical replicates. “F”; indicator for analysis strategy with “−”; no FAIMS, “10”; 10 Th window width and 20 ms DIA injection time, “20”; 20 Th window width and 40 ms DIA injection time. (B) As **A**, but data analysis using Spectronaut 19, with gray and blue corresponding to data analysis using Method Evaluation mode and directDIA mode, respectively. (C) Total extracted ion current (XIC) profile across the chromatographic gradient, for a representative 1 pg run without FAIMS (top) or when using FAIMS (bottom). Ion current was stratified by charge state 1 to 3, with unknown or 4+ charge states omitted from the visualization. The plot on the right side is a zoomed-in version of the FAIMS run, highlighting the almost total absence of atmospheric contamination (*z* = 1). (D) Dilution series experiment as described in **A**, but using FAIMS in all cases and comparing the difference between the Orbitrap Astral and Orbitrap Astral Zoom using a prototype mass spectrometer. Percentage increases of median values are indicated, * = *p* < 0.05 via two-tailed Student’s t-testing, *n* = 4 technical replicates. (E) As **D**, but data analysis using Spectronaut 19, with gray and blue corresponding to data analysis using Method Evaluation mode and directDIA mode, respectively. Differences of medians were not assessed in directDIA mode due to the non-linear behavior of the directDIA identification strategy. (F) Dilution series experiment as described in **A**, injecting between 50 and 2,000 pg of HeLa lysate on a 50 SPD gradient, and using MS and FAIMS methodology optimized for MS1-based quantification and for each sample load. Number of identified protein-coding genes are shown for data analyses using either DIA-NN or Spectronaut. Bars represent median, *n* = 3 technical replicates. (G) As **F**, but showing number of precursors (DIA-NN) or stripped peptide sequences (Spectronaut). (H) Violin plot distribution of coefficient of variation for all protein-coding genes from **F**. MS1-based quantification via Spectronaut, MS2-based quantification via DIA-NN. Thick line represents median, thin lines 1^st^ and 3^rd^ quartiles.

Next, we investigated the gain in performance across the dilution series using our Orbitrap Astral Zoom MS prototype, comparing regular Astral mode versus Astral Zoom mode with all its features enabled (Fig. 3D-E and S6C-H). In the range from 10 to 100 pg, we observed consistent increases of 5-9% at the precursor level and 2-4% at the protein level when using DIA-NN in first pass mode (Fig. 3D and S6C). In the 1 to 5 pg range, precursors increased by 9-12% and proteins increased by 11-15%. At the lowest quantified load of 0.5 pg, increases in the range 40-60% were observed; however, these occurred at the very limit of detection. Similar increases were found with Spectronaut data analysis (Fig. 3E and S6D), although more sporadically depending on input amount. Quantitatively, when inspecting coefficient of variation of the MS2-level quantification, we found that the Astral Zoom was able to achieve similar coefficient of variation with approximately half the input amount as compared to the Astral (Fig. S6E-F). Pearson correlations between same-load replicates were comparable or slightly enhanced with the Astral Zoom (Figure S6G-H).

Quantitative precision and reproducibility are important tenets of proteomics method design. We have primarily performed MS2-based quantification throughout this study; however, a core strength of the Astral and Astral Zoom is that they can dedicate the Orbitrap mass analyzer to MS1-based quantification without sacrificing the ability to sequence precursors in parallel using the Astral mass analyzer (Guzman et al., 2024). To exemplify the capability of the Astral Zoom to leverage MS1-based quantification, we performed another dilution series experiment ranging from 2,000 pg down to 50 pg of HeLa, using 50SPD gradients and FAIMS (Fig. 3F-G). From 50 pg, we could identify ∼5,100 proteins using DIA-NN in first pass mode, and up to ∼7,200 proteins when co-analyzing the 50 pg runs together with all other loads using Spectronaut in directDIA mode. Importantly, using MS1-based quantification, we achieved exceptionally low median coefficients of variation of 5.2-6.5% at the protein level and 10.4%-11.8% at the precursor level (Fig. 3H and S6I).

Overall, our data showed that the combination of FAIMS and the Astral Zoom enables reproducible and accurate proteome profiling from protein inputs far below that of a typical single HeLa cell, paving the way for analysis of smaller or less protein-rich cells.

### Optimization of the CellenOne-based single-cell proteomics workflow

Having optimized and benchmarked our LC-MS methodology using a diluted HeLa lysate, we next sought to adapt these advances to the analysis of individual, physically sorted and isolated HeLa cells. Single-cell proteomics introduces unique experimental challenges, including limited material, surface adsorption, and variability in lysis and digestion efficiency. To address these, we established and evaluated a single-cell workflow using the CellenOne for cell isolation and digestion (Fig. 4A). Notably, for all SCP experiments in this study, we did not perform peptide clean-up, and digested single cells were directly injected onto the analytical LC column for online LC-MS analysis.

**Figure 4.**
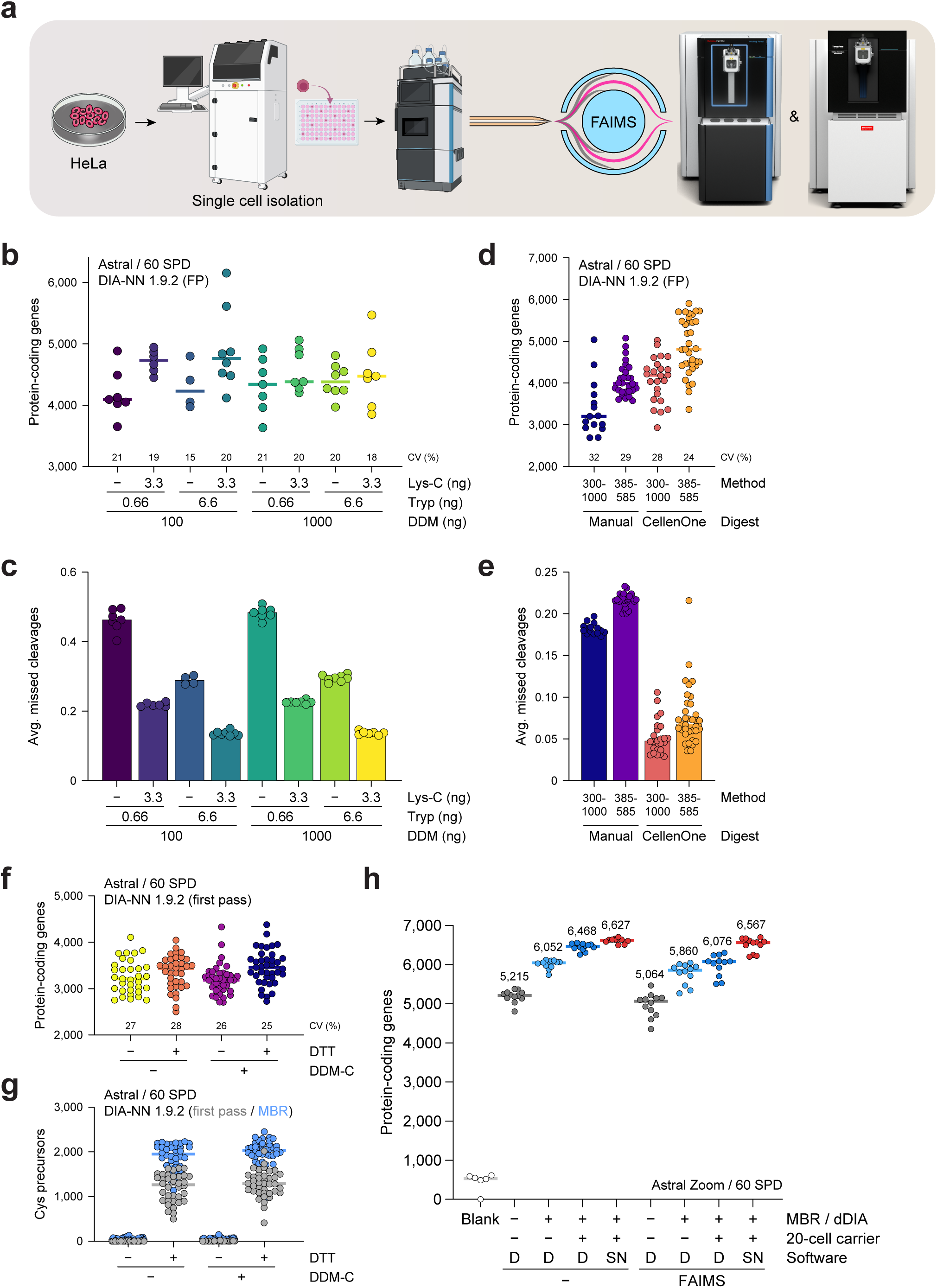
Optimization of single-cell proteomics MS methodology for standard and FAIMS workflows. (A) Schematic overview of the experimental design. HeLa cells were grown, single cells were sorted into and digested in 96-well plates using the CellenOne platform, and digests were directly analyzed using a Vanquish HPLC, laser-pulled and in-house packed analytical columns, and an Orbitrap Astral Zoom mass spectrometer prototype. (B) Number of protein-coding genes identified from single HeLa cells when using various different digestion mixes. Cells were sorted by the CellenOne platform, but otherwise manually digested. Bars represent median. Numbers above the x-axis are median coefficient of variation. *n* = 4-8 single cells. (C) As **B**, but showing the average number of missed tryptic cleavages observed per peptide. (D) As **B**, but comparing manual digestions (with the CellenOne only used to dispense single cells) to fully automated handling and digestion using the CellenOne, as well as comparing the broader-scan range nDIA method with ultra-sensitive narrow-scan range nDIA. *n* = 15-35 single cells. (E) As **D**, but showing the average number of missed tryptic cleavages observed per peptide. (F) As **B**, but additionally comparing the effect of dithiothreitol (DTT) in the digestion mix, as well as pre-coating the 96-well plate with n-dodecyl-ß-D-maltoside (DDM). *n* = 33-40 single cells. (G) As **F**, but showing the number of cysteine-containing precursors identified in the data. (H) Number of protein-coding genes identified from single HeLa cells (median diameter ∼26.5 μm) when using optimized sample preparation workflows, computational inclusion of 20-cell carriers, and using either standard-or FAIMS-optimized methodology. Data analysis using either DIA-NN (“D”) or Spectronaut (“SN”) as indicated. *n* = 12 single cells. “MBR”; DIA-NN matching-between-runs, “dDIA”; Spectronaut directDIA.

We first performed an evaluation of different compositions of the combined cell lysis, protein extraction, and digestion buffer (e.g. master mix), wherein we varied concentrations of DDM, trypsin, and Lys-C (Fig. 4B-C and S7A-C). For this experiment, we manually performed deposition of master mix and digestion, whereas single cell isolation into the wells was performed using a CellenOne. Overall, we were able to identify between 4,000 and 5,000 proteins per cell using DIA-NN in first pass mode (Fig. 4B), and up to ∼5,500 proteins per cell when using match-between-run (MBR) mode (Fig. S7B), regardless of master mix composition, although there was a trend for higher identification rates when both Lys-C and trypsin were present in the master mix. Notably, when considering missed tryptic cleavage rate (Fig. 4C), we observed a very consistent digestion efficiency for all cells digested with the same master mix, with the Lys-C and trypsin combination necessary for achieving the lowest fraction of missed tryptic cleavage rates.

Next, we compared manual dispensing of master mix and digestion with fully automated processing using the CellenOne, which was used for single-cell isolation in both cases (Fig. 4D-E and S7D-H). For each of the approaches, we analyzed single cells using either regular nDIA or ultra-sensitive nDIA methods. Consistent with what we observed with diluted HeLa lysate (Fig. 1F), we identified fewer precursors, but considerably more (∼15-25%) proteins when using the ultra-sensitive nDIA method (Fig. 4D). Strikingly, using the CellenOne for all steps of the single-cell processing dramatically reduced the number of missed cleavages from ∼20% to ∼5% (Fig. 4E), with a concomitant increase of 20-30% more proteins identified per cell (Fig. 4D). With computational inclusion of a 20-cell carrier and using DIA-NN in MBR mode, we were able to identify a median of ∼6,000 proteins per cell (Fig. S7G).

Cysteine disulfide bridge reduction and alkylation of proteins is not routinely performed in SCP workflows (Bubis et al., 2025; Ye et al., 2025), due to the challenges associated with sample loss, introduction of chemical contaminants, and the potential for off-target chemical modifications when handling minute amounts of protein. As our workflow does not include a peptide clean-up step, we were limited to using only volatile or otherwise MS-compatible reagents. We reasoned that reduction during single-cell protein digestion using dithiothreitol (DTT) would be sufficient to reduce cysteine residues until stabilization via acidification of the digest. We performed a manual master mix dispensing and single-cell digestion step, wherein we compared the absence or presence of DTT, as well as the effect of coating the 96-well plate in DDM prior to master mix deposition (Fig. 4F-G and S8A-G). Coating the 96-well plate with DDM had no discernable effect, likely because the master mix itself also contained DDM. Addition of DTT resulted in an ∼8% increase at the protein level (Fig. 4F), and an ∼18% increase at the precursor level (Fig. S8A). Expectedly, in the absence of DTT, we could not detect any cysteine-containing precursors, whereas up to ∼2,000 cysteine-containing peptide precursors were readily detectable with DTT (Fig. 4G).

Finally, we compared analyzing single HeLa cells on the Astral Zoom using either standard or FAIMS acquisition (Fig. 4H and S8H-I). Overall, we identified ∼10% fewer precursors when using FAIMS (Fig. S8H), but a comparable number of proteins (Fig. 4H), with ∼5,100 proteins per cell when using DIA-NN in first pass mode, and ∼6,500 proteins per cell when including 20-cell carriers and using either DIA-NN in MBR mode or Spectronaut in directDIA mode. Coefficients of variation were not notably different between standard and FAIMS methods (Fig. S8I). Using blank injections between single cell runs, only a few hundred proteins were identified, indicating a run-to-run carryover of ∼0.4% (Fig. S8J). We also performed an investigation of the computational proteome carrier effect via co-analysis of single or several HEK cells, as described in Supplementary Note 3.

Taken together, our results highlight key parameters that govern single-cell proteomics performance, including enzyme composition, reduction conditions, and automation of sample handling. The resulting CellenOne-based workflow delivers high digestion efficiency, minimal carryover, and proteome coverage exceeding 6,000 proteins per cell at a throughput of 60 SPD, underscoring its suitability for sensitive and reproducible single-cell proteomics.

### Ultra-deep profiling of single HeLa cells uncovers low-abundant proteins

Having established an efficient workflow for isolation and digestion of single cells, as well as enhanced MS methodology for analysis of low input amounts, our next goal was to achieve comprehensive depth of sequencing within single HeLa cells while also maintaining good quantitative precision. For this, we performed another series of single HeLa experiments based on our optimized CellenOne-based protocol. For the SCP LC-MS analysis, we utilized FAIMS in combination with extended gradients (40 SPD), bolstered by the enhanced sensitivity of the Orbitrap Astral Zoom MS. From single HeLa cells and without inclusion of carriers, we were able to identify ∼56,000 or ∼69,000 peptides, and ∼6,600 or ∼7,100 proteins, respectively with DIA-NN or Spectronaut (Fig. 5A-B). Computational inclusion of carriers was able to boost the number to ∼7,300 proteins from single HeLa cells. Considering our depth of sequencing, we performed a database search for phosphorylation as a variable modification on serine, threonine and tyrosine residues (Bekker-Jensen et al., 2020), and were able to find a median of 230 localized phosphorylation sites in single HeLa cells (Fig. 5A), which is about twice as high as observed previously (Ye et al., 2025).

**Figure 5.**
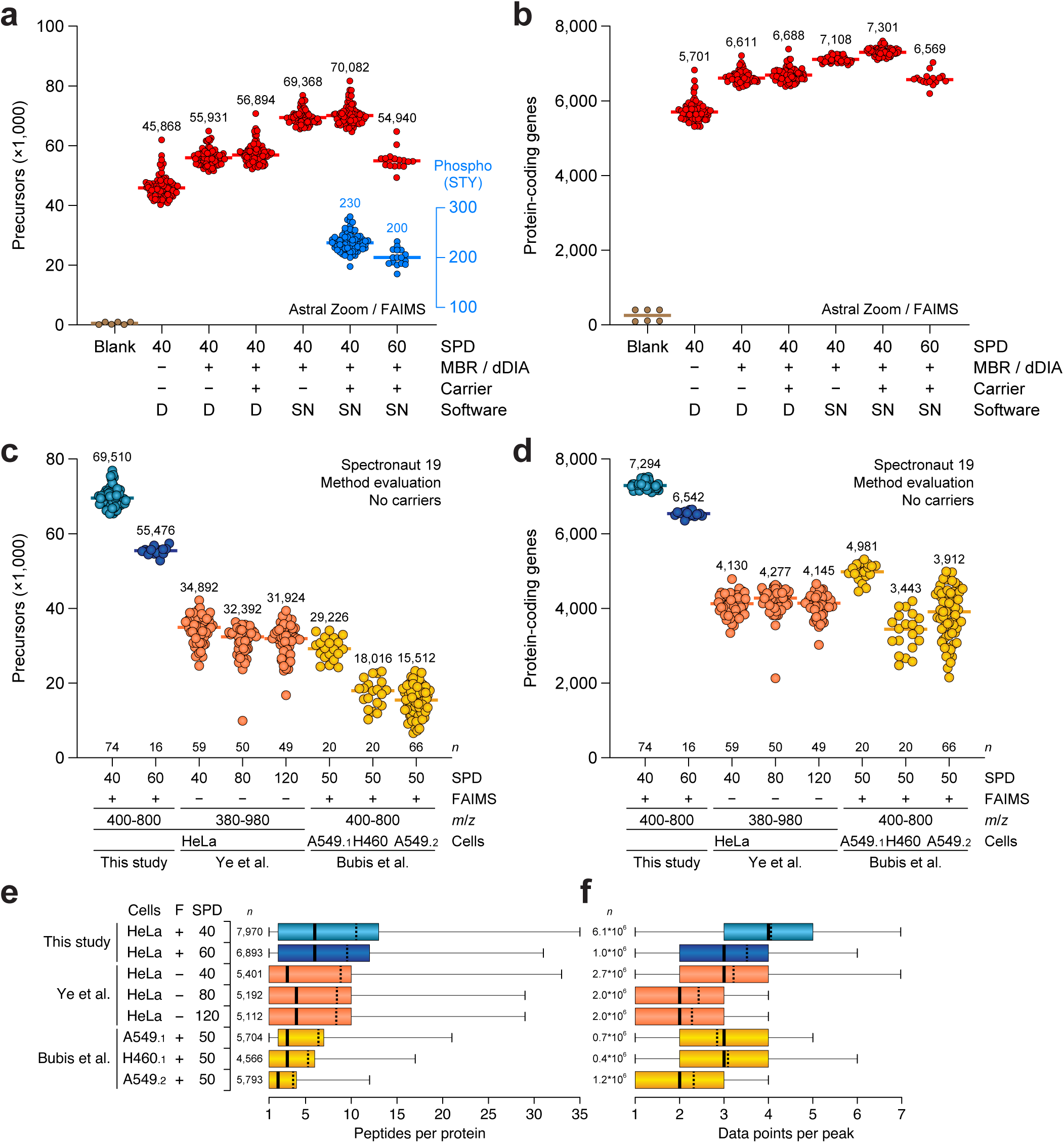
Robust profiling of >7,000 proteins from single HeLa cells. (A) In red, number of precursors (DIA-NN) or stripped peptide sequences (Spectronaut) identified from single HeLa cells (median diameter ∼26.5 μm). In blue, number of localized (>0.75 probability) phosphorylation sites identified from single HeLa cells. Automated single cell isolation and digestion using the CellenOne, all analysis on Orbitrap Astral Zoom MS with FAIMS, comparing 40 and 60 samples-per-day (SPD) gradients. Bars represent median values, exact medians displayed above each group of data. Data analysis using DIA-NN (“D”) or Spectronaut (“SN”) as indicated. “Carrier”; computational carrier of 20, 40, and 60 cells. “MBR”; DIA-NN matching-between-runs, “dDIA”; Spectronaut directDIA. *n* = 74 single cells (for 40 SPD) and *n* = 16 single cells (for 60 SPD). (B) As **A**, but for the number of protein-coding genes. (C) Comparison between the number of precursors identified from single cells in this study, to two recent landmark papers using similar single-cell approaches (Bubis et al., 2025; Ye et al., 2025). Publicly available raw MS data files were downloaded and re-processed in one computational run using Spectronaut in Method Evaluation mode. Bars represent median values, exact medians displayed above each group of data. Number of replicate single cells (*n*) is displayed above the x-axis. Appendix numbers following cell type indicate two separate batches of A549 cells from Bubis et al. (D) As **C**, but for protein-coding genes. (E) As **D**, quantifying the number of peptides identified per protein-coding gene, visualized as a box plot. Thick line; median, dashed line; average, box limits; 1^st^ and 3^rd^ quantile, whisker limits; 5^th^ and 95^th^ percentile. *n* = the total number of cumulative protein-coding genes identified within each cell group, as indicated. (F) As **E**, but visualizing the number of data points per peptide peak, with higher numbers of data points allowing more precise quantification. *n* = the total number of traceable peaks within the entire cell group, as indicated.

Two landmark single-cell proteomics studies were recently published, which used comparable approaches and MS instrumentation applied to human epithelial cancer cell lines like HeLa and A549 (Bubis et al., 2025; Ye et al., 2025). To benchmark our optimized SCP workflow described here, we performed a direct comparison of our SCP data to theirs, via re-analysis of our MS data together with their publicly available raw MS data. We searched all data using the same settings to minimize computational variance, and we excluded carrier channels, as the numbers and types of carriers were highly irregular between studies. Reassuringly, we found similar numbers of precursors and proteins from our own data even when searched with slightly different settings and together with data from other studies (Fig. 5C-D and S9A-D). From our 40 SPD runs, we identified two-to three-fold more precursors, and 2,000-3,000 more proteins from single cells compared to the two landmark studies. Notably, we also achieved greater numbers of identifications from our 60 SPD runs. To assess quantitative capability of our method, we investigated peptides-per-protein as well as data points per peak, and compared this across studies. With both our 40 and 60 SPD methods, we achieved a median of six peptides per protein, whereas other studies only identified two to four peptides per protein (Fig. 5E). In terms of data points per peak, we achieved a median of three to four points per peak for 60 and 40 SPD, respectively, with other studies generally getting two to three points per peak (Fig. 5F). Together, this demonstrates that both qualitatively and quantitatively we achieve considerably greater numbers compared to contemporary methodology.

To evaluate the biological implications of our deep profiling of single HeLa cells, we examined which specific proteins we identified in comparison to the two other studies (Bubis et al., 2025; Ye et al., 2025). To this end, we performed a Venn comparison between all three datasets, and also an UpSet analysis to highlight overlapping sets of proteins across all different conditions (Fig. 6A and S10A). Expectedly, when considering proteins identified by at least two peptides and in at least half of single cells (Fig. 6A), the largest overlapping set of proteins was shared among all studies (3,285 proteins), and even across all conditions (1,699 proteins). The second- and third-largest intersections were represented by proteins exclusively identified in this study, followed by a mixture of intersections dependent on study size and technical parameters. Next, we evaluated the depth of sequencing achieved by us and others (Fig. 6B and S10B), through mapping all identified proteins to copy-numbers determined in an ultra-deep HeLa proteome study (Bekker-Jensen et al., 2017). As anticipated, all SCP studies identified essentially all of the most abundant proteins, regardless of technical conditions and cell type. Strikingly, we were able to identify and quantify proteins with significantly lower copy-numbers compared to other studies, in some cases detecting three to five times more low-abundant proteins (Fig. 6B). When considering all proteins, even those identified by a single peptide, largely the same patterns were observed (Fig. S10). Overall, we were able to identify 1,988 proteins of below-average abundance that were not consistently identified by other contemporary SCP studies. Intriguingly, when inspecting Uniprot keywords associated with these proteins through term enrichment analysis, we found a significant overrepresentation of biologically relevant terms such as DNA repair, phosphoproteins, chromatin regulation, cell cycle, and proteins with nuclear and chromosomal localization (Fig. 6C).

**Figure 6.**
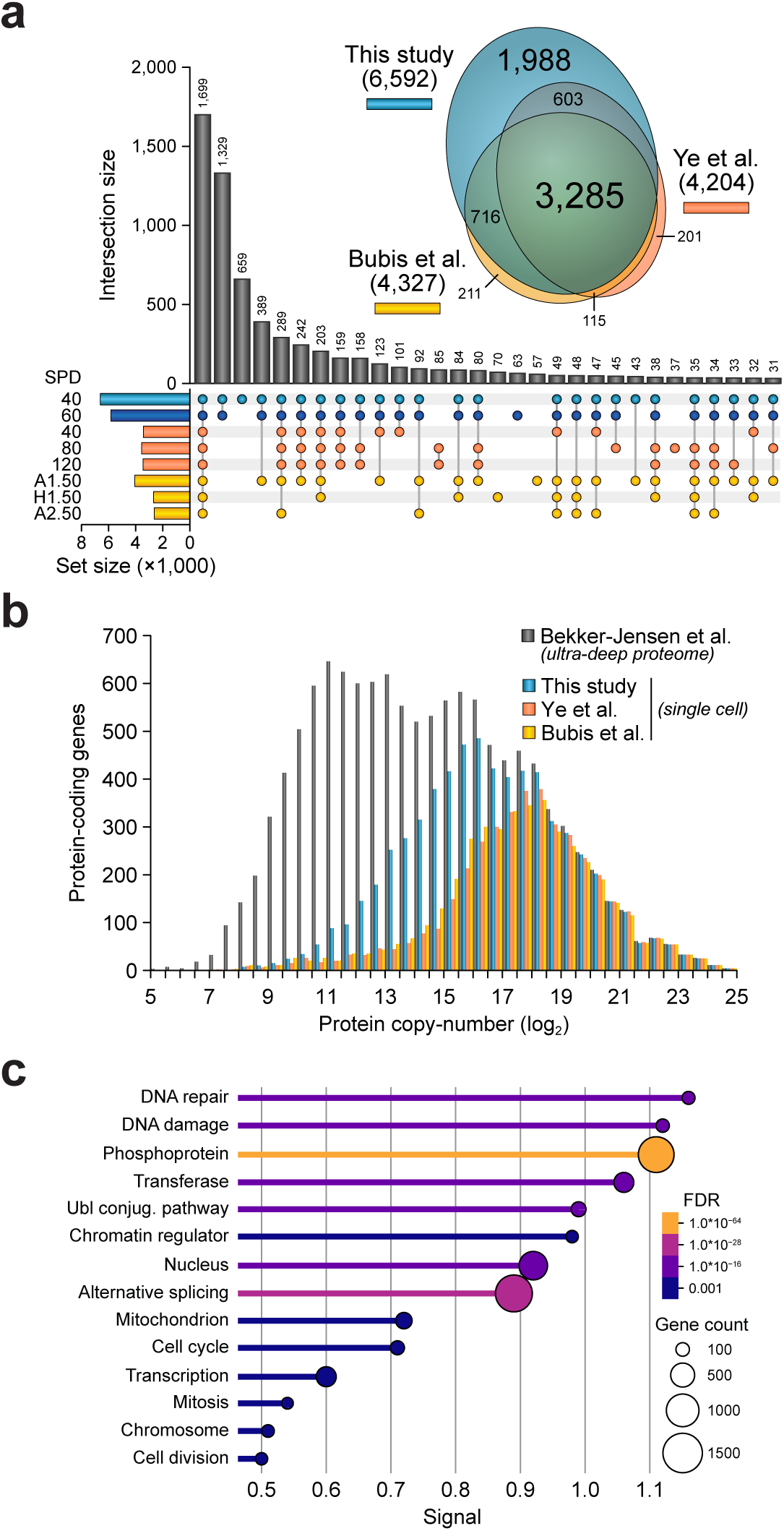
Greatly enhanced single cell proteome depth compared to contemporary single cell studies. (A) UpSet plot analysis, visualizing distinct and overlapping sets of proteins identified from single HeLa cells in this study compared to two recent studies using similar single-cell approaches (Bubis et al., 2025; Ye et al., 2025), see also Figure 5C-F. Exact values are displayed above each bar. Analysis based on re-processing of available MS data using Spectronaut in Method Evaluation mode, only proteins identified by 2 or more peptides and detected in at least 50% of replicates were considered. All single cells were HeLa, except for “A1”; A549 batch 1, “H1”; H460 batch 1, “A1”; A549 batch 2. The in-set scaled Venn diagram depicts global overlap between the three studies. (B) Depth of sequencing visualization, with known protein copy-numbers taken from a published ultra-deep proteome. Proteins identified in all single-cell studies were mapped to the total proteome and binned in 0.5 log_2_ bins. (C) Uniprot keyword enrichment analysis for the 1,988 protein-coding genes uniquely identified in this study (see **A**), querying generic Uniprot keywords via the STRING database (Szklarczyk et al., 2023). “Signal” is an admixture of both term enrichment and FDR, as reported by the STRING database. All top terms and several other representative (non-duplicate) terms are shown, for the full list of enriched terms refer to Supplementary Table Y. FDR and protein-coding gene count in for each term are indicated.

Collectively, this work pushes the frontier of single-cell proteomics, achieving unparalleled coverage and precision from individual HeLa cells. By integrating enhanced LC-MS gradients and methodology, FAIMS, and high-sensitivity Orbitrap Astral Zoom MS detection, we resolve >7,000 proteins per single HeLa cell, including ∼2,000 previously undetectable lower copy-number proteins. Our approach not only redefines the analytical depth achievable in single-cell studies but also opens new avenues for probing the regulatory proteome underlying cell identity and function.

### Ultra-sensitive SCP captures the immune landscape of PBMCs

Small cells below a certain size (< ∼10 µm in diameter) are frequently excluded from SCP analyses. This is because their low protein content leads to insufficient proteome coverage, a problem that is exacerbated by the cubic relationship between cell diameter and protein yield, where small decreases in diameter lead to disproportionately large drops in detectable protein. This has made robust proteome analysis technically unfeasible for very small cells, and their inclusion can introduce significant noise and missing data into SCP datasets.

Peripheral blood mononuclear cells (PBMCs) are commonly investigated in clinical practice (Franciosa et al., 2023; Krieg et al., 2018), because they represent a minimally invasive and accessible window into systemic immune activity. The PBMC population consists primarily of lymphocytes (T cells, B cells, and natural killer cells) (Kleiveland, 2015), and the majority are very small cells (6-8 µm in diameter) (Kuse et al., 1985). Contemporary study of PBMCs relies on bulk analysis, where billions of cells are co-analysed and cell-to-cell heterogeneity is lost (Koncarevic et al., 2014; Santa et al., 2024). Here, to evaluate and benchmark our enhanced SCP methodology in the context of some of the smallest cells, we applied it to PBMCs directly derived from human blood (Fig. 7A). To visualize the difference in size between PBMCs and human epithelial cancer cells, e.g HeLa and HEK293T cells, we plotted the size distribution of all single cells isolated in this study (Fig. 7B and S11A). HEK293T and HeLa cells are generally in the range of 20 to 30 µm, whereas a mixed population of PBMCs contained cells in the range of 6 µm to 16 µm, with the smallest 6-8 µm cells being the most numerous, corresponding to ∼50 times less protein per cell.

**Figure 7.**
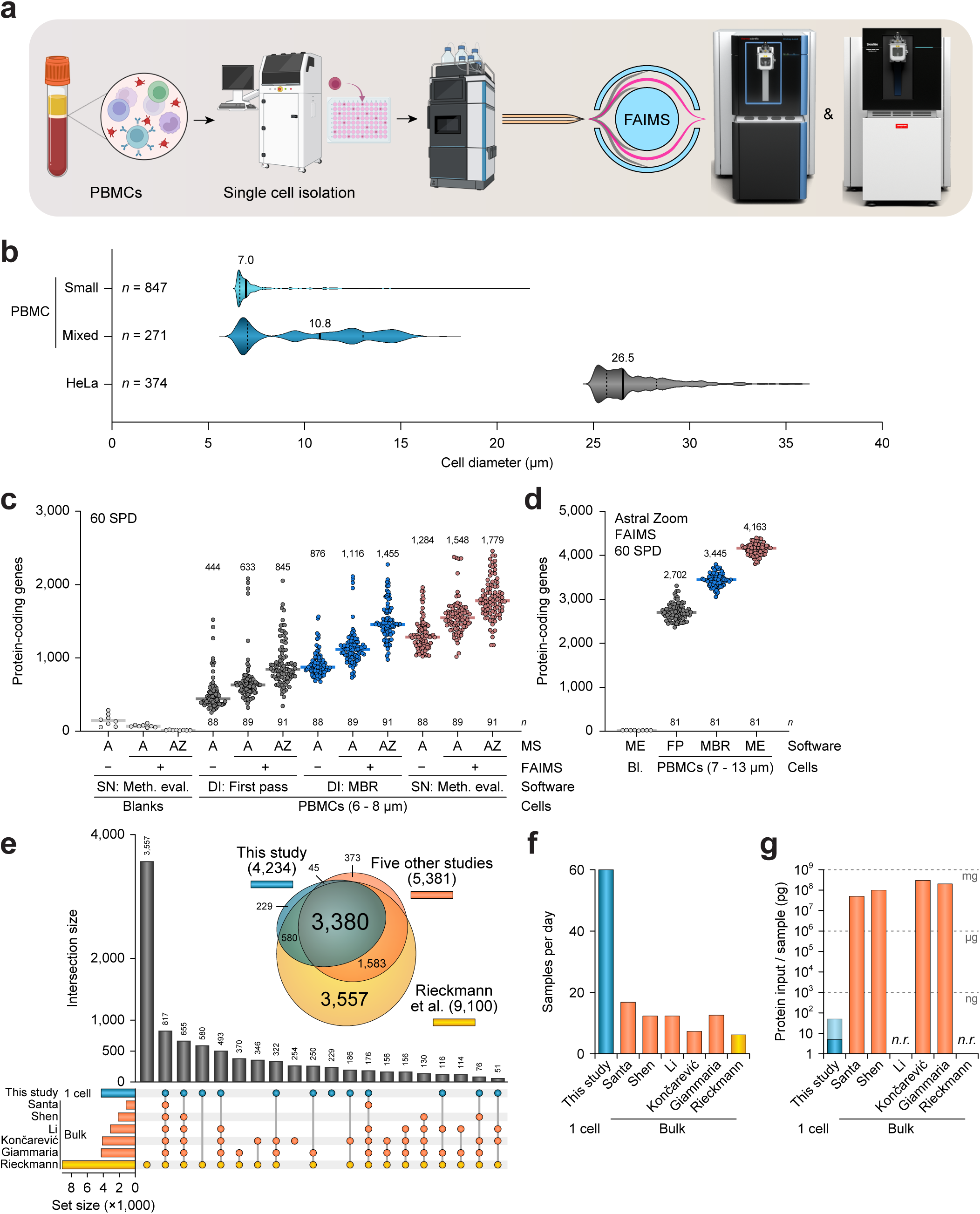
Profiling single peripheral blood mononuclear cells with proteome depth comparable to bulk analysis. (A) Schematic overview of the experimental design. Peripheral blood mononuclear cells (PBMCs) were isolated from buffy coats derived from anonymous donor blood, and single PBMCs were sorted into and digested in 96-well plates using the CellenOne platform, and digests were directly analyzed using a Vanquish HPLC, laser-pulled and in-house packed analytical columns, and an Orbitrap Astral Zoom mass spectrometer prototype. (B) Size distribution profile of all PBMCs isolated in this study. PBMCs were prepared to exclusively yield the smallest cells (“Small”), or minimally processed to gain a mixed population (“Mixed”). HeLa cells are shown for reference (see Figures 4-5). Thick lines represent median, with median values indicated, and thin dashed lines indicate 1st and 3rd quartiles. *n* = number of sorted cells as indicated. (C) Number of protein-coding genes identified from the smallest PBMCs (median diameter ∼7 μm), comparing performance of an Orbitrap Astral (“A”) with and without FAIMS, to an Orbitrap Astral Zoom (“AZ”) with FAIMS. Bars represent median values, exact medians displayed above each group of data. Data analysis using DIA-NN (“DI”) or Spectronaut (“SN”) as indicated. “MBR”; DIA-NN matching-between-runs, “dDIA”; Spectronaut directDIA. *n* = 88-91 single PBMCs. (D) As **C**, but visualizing analysis of the Mixed PBMC population using the Orbitrap Astral Zoom with FAIMS. *n* = 81 single PBMCs. “ME”; Spectronaut with Method Evaluation, “FP”; DIA-NN with first pass, “MBR”; DIA-NN with matching-between-runs. (E) UpSet plot analysis, visualizing distinct and overlapping sets of proteins identified in this study from single PBMCs, to proteins identified by published bulk PBMC studies using a wide range of different technical approaches (Giammaria et al., 2025; Koncarevic et al., 2014; Li et al., 2022; Rieckmann et al., 2017; Santa et al., 2024; Shen et al., 2019). To ensure stringency, proteins identified from single PBMCs in this study were only considered if identified by both DIA-NN and Spectronaut. Proteins from other studies were taken as reported by the respective authors, with data filtering and alignment steps described in methods and Supplementary Table Y. Exact values are displayed above each bar. The in-set scaled Venn diagram depicts global overlap between proteins identified from single PBMCs in this study, to the summation of five intermediate-sized bulk PBMC studies (“Five other studies”), and proteins reported by an in-depth and comprehensive bulk PBMC profiling by Rieckmann et al. (F) As **E**, indicating the sample throughput of the methodology used in each study. (G) As **E**, visualizing the amount of starting material in picograms per sample, for each study. Lower numbers indicate reduced material requirement, and are thus favorable. “n.r”; not reported.

First, we performed an investigation of the smallest PBMCs, where we processed the PBMCs to exclude larger cells prior to CellenOne isolation and digestion (Fig. 7B). In this experiment, we compared data acquisition using the Astral without FAIMS, the Astral with FAIMS, and the Astral Zoom with FAIMS (Fig. 7C and S11B). Generally, we were able to identify ∼4,000 to ∼6,500 precursors per single PBMC, and ∼900 to ∼1,800 proteins per single PBMC. In all cases, FAIMS notably enhanced identifications at both the precursor and protein levels (+21% to +43%), consistent with what we observed in our dilution series experiments at ultra-low input amounts (Fig. 3A-B and S6A-B). Furthermore, using the Orbitrap Astral Zoom MS, we could profile more proteins (+15% to +33%), similarly in line with what we found previously (Fig. 3D-E and S6C-D). Next, we investigated the mixed population of PBMCs that were minimally processed prior to CellenOne isolation and digestion, with all data acquisition performed using the Astral Zoom with FAIMS (Fig. 7D and S11C). From this experiment, we could identify ∼19,000 to ∼26,000 precursors and ∼3,400 and ∼4,200 proteins from single PBMCs, respectively with DIA-NN and Spectronaut.

To validate our single PBMC data, we performed a comprehensive comparison against bulk PBMC proteomics studies (Giammaria et al., 2025; Koncarevic et al., 2014; Li et al., 2022; Rieckmann et al., 2017; Santa et al., 2024; Shen et al., 2019), which currently represent the vast majority of MS-based PBMC investigation. To this end, we overlapped proteins identified from single PBMCs in this study to six published bulk PBMC proteomics studies that utilized a wide range of experimental designs and LC-MS instrumentation, and performed both Venn diagram comparison and UpSet plot analysis to highlight overlapping sets of proteins across all studies (Fig. 7E and S11D). Strikingly, compared to the majority of bulk PBMC studies, we identified as many or more proteins from single PBMCs. A notable exception was the human hematopoietic cell atlas study by Rieckmann et al., which mapped significantly more proteins through in-depth profiling of 28 primary human hematopoietic cell populations in steady and activated states (Rieckmann et al., 2017). Otherwise, the second-largest intersection represented 817 proteins identified across all 7 studies, and generally we observed a strong overlap between proteins identified in this work from single PBMCs versus proteins identified from bulk PBMC Fig. 7E and S11D). Importantly, we analyzed our samples at considerably higher throughput (Fig. 7F), and used seven orders of magnitude (∼10 million) times less input material compared to bulk PBMC studies (Fig. 7G), demonstrating the exquisite sensitivity of our methodology.

From a biological perspective, when considering the tissue expression database TISSUES localization annotation (Palasca et al., 2018), the proteins we identified from single PBMCs were functionally overrepresented for blood-related terms, including blood plasma, lymphocyte, leukocyte, and the immune system (Fig. 8A). When considering Reactome annotation (Milacic et al., 2024), the largest clusters of proteins we identified from single PBMCs were associated with the immune system, infectious disease, and the cellular response to stress (Fig. 8B). In relation to bulk PBMC studies (Giammaria et al., 2025; Koncarevic et al., 2014; Li et al., 2022; Rieckmann et al., 2017; Santa et al., 2024; Shen et al., 2019), we found as many or more proteins related to the innate immune system, adaptive immune system, and infectious disease (Fig. 8C), and in similar distributions. Although we did not profile as many immune-related proteins as Rieckmann et al., our data showed a more significant statistical overrepresentation (Fig. S12A). Within the adaptive immune response, we noted significant coverage over five out of eight pathways annotated by Reactome (Fig. S12B), including T cell regulation and signaling, B cell signaling, and MHC Class I and II antigen presentation. Among overrepresented KEGG pathways, one notable example also included the T cell receptor signaling pathway, with a significant coverage over ∼55% of proteins involved in the pathway (Fig. 8D), in line with ∼61% coverage over the same pathway in Reactome (Fig. S12B).

**Figure 8.**
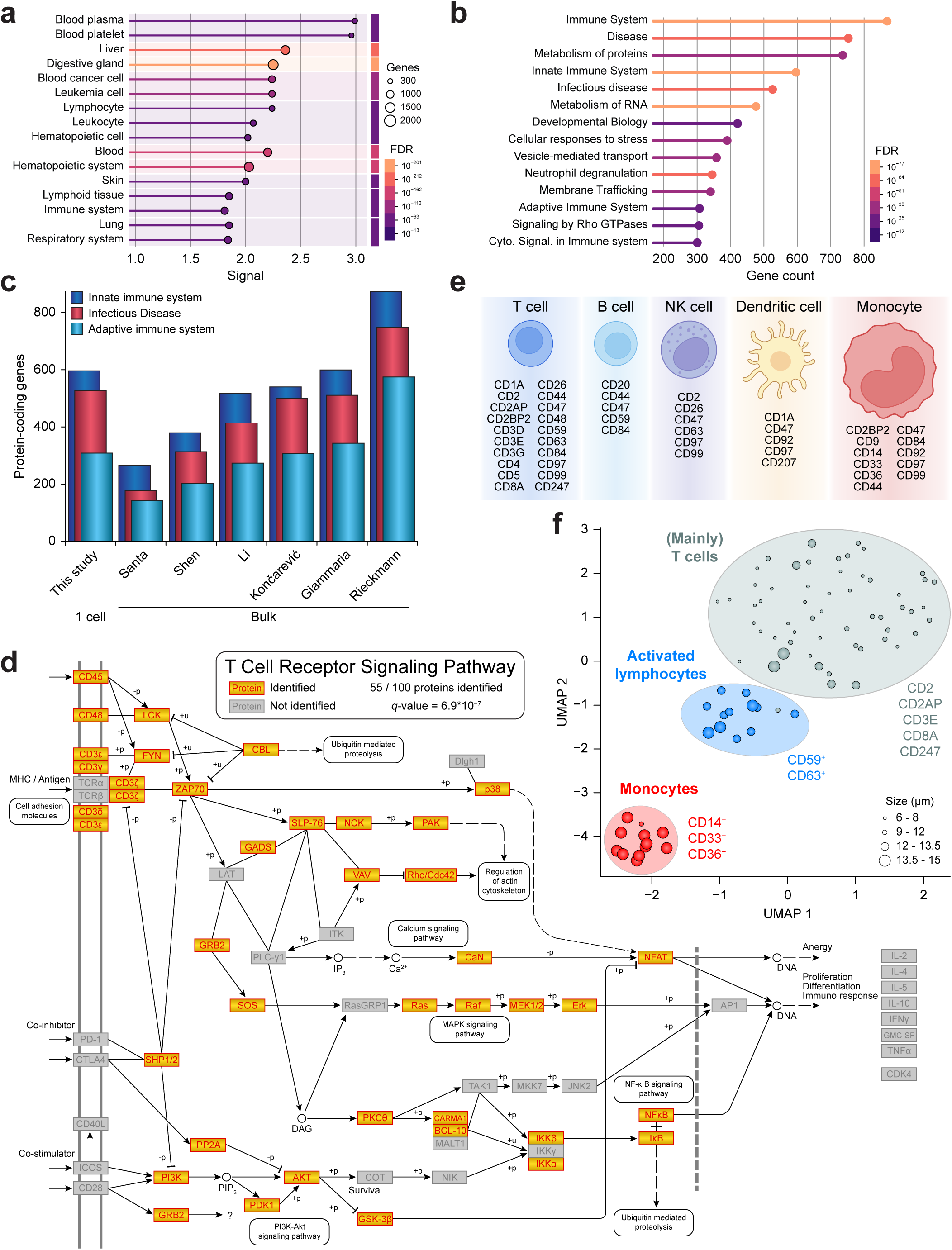
Insights into distinct PBMC cell populations at the single cell level. (A) TISSUES term enrichment analysis on all protein-coding genes identified from single PBMCs in this study (see Figure 7), querying for TISSUES annotation via the STRING database (Szklarczyk et al., 2023). “Signal” is an admixture of both term enrichment and FDR, as reported by the STRING database. The top 10 clusters of over-representation terms are shown, for the full list of enriched terms refer to Supplementary Table Y. FDR and protein-coding gene count in for each term are indicated. (B) As **A**, but for Reactome Pathway enrichment. The largest enriched categories are visualized, for the full list of enriched terms refer to Supplementary Table Y. FDR for each term is indicated. (C) As **B**, but visualizing the number of protein-coding genes corresponding to the indicated immune-related Reactome categories, as identified in this study from single PBMCs, or from bulk PBMC in six other studies (Giammaria et al., 2025; Koncarevic et al., 2014; Li et al., 2022; Rieckmann et al., 2017; Santa et al., 2024; Shen et al., 2019). (D) Kyoto Encyclopedia of Genes and Genomes (KEGG) pathway analysis (Kanehisa and Goto, 2000), mapping all protein-coding genes identified in this study from single PBMCs to the “T Cell Receptor Signaling Pathway” as curated by KEGG. Overall protein identification number and the accompanying FDR value were derived through the STRING database (Supplementary Table Y). (E) Overview of all Cluster of Differentiation (CD) and associated proteins identified from single PBMCs, grouped by the class of PBMC these markers are usually observed in, based on Rieckmann et al (Rieckmann et al., 2017). (F) Euclidian UMAP analysis of the Mixed PBMC population, using all CD proteins identified by DIA-NN and Spectronaut to drive the clustering. Orthogonal unsupervised hierarchical Pearson clustering was used to trace the two most distinct cellular clusters of CD markers (Figure **S12C**), which are colored red and blue, with the over-represented CD proteins indicated in the same colors. Remaining cells are colored in gray, with various small clusters of heterologously observed CD markers indicated. Sizes of all individual cells, as recorded by the CellenOne during sorting, are indicated. *n* = 81 single PBMCs.

As PBMCs represent a heterogeneous mixture of different cell types, we wondered whether we could distinguish between cell types exclusively based on our single PBMC proteomics data, without any pre-sorting or other orthogonal techniques to provide cellular identity. To this end, we investigated all Cluster of Differentiation (CD) proteins present in our single PBMC data, as these proteins strongly correlated to cellular identity and are routinely used distinguish between cell types (Engel et al., 2015). Across our PBMC experiments, we could reliably detect 26 CD proteins (Fig. 8E), which we assigned to the five main classes of PBMC based on the quantitative data from Rieckmann et al. We observed most markers corresponding to T cells, followed by monocytes, although in general CD marker proteins are not always exclusive to one single cell type. Finally, utilizing only the CD proteins identified from our single PBMC analysis, we performed a Euclidian UMAP clustering analysis (Fig. 8F). We additionally performed unsupervised hierarchical Pearson clustering to find overrepresented clusters of CD proteins (Fig. S12C), through which the two most notable clusters were overrepresented in either CD14/33/36 or CD59/63. When superimposing these clusters onto the UMAP, along with cellular sizes as determined during single-cell sorting, we found two distinct clusters of cells corresponding to monocytes and activated lymphocytes (Fig. 8F), highlighting the strength of our strategy by demonstrating that we are able to distinguish distinct cell types from a mixed population without prior knowledge.

By pushing SCP sensitivity to its limits, we achieved comprehensive proteome profiling of individual PBMCs – immune cells nearly 100 times smaller in volume than HeLa cells – at throughput and depth rivaling bulk studies. Our approach uncovered thousands of immune-associated proteins, captured comprehensive functional pathway coverage, and resolved distinct immune cell populations based purely on proteomic signatures. These findings establish a foundation for high-resolution mapping of immune heterogeneity directly from patient-derived samples.

## DISCUSSION

In this study, we systematically optimized and integrated each component of the single-cell proteomics (SCP) workflow, from peptide recovery and chromatographic design to data acquisition and instrument configuration. Our investigation of chromatography and surfactant-assisted recovery establishes a cohesive low-input workflow that improves both depth and throughput from low-picogram samples. We find that presence of DDM in the lowest amount we tested (∼1,000 fold weight excess over peptide) was sufficient to increase identification rates from low-input samples, with higher amounts of DDM providing no further benefit. Indeed, DDM is routinely applied in the context of SCP to assist in peptide recovery and cell lysis (Tsai et al., 2021). Although concentrations of 0.1% DDM are typically used in master mix, we did not observe a difference in protein recovery efficiency when going down to 0.01% DDM, which could suggest that the beneficial effect of DDM primarily relates to peptide recovery, rather than cell lysis. On the chromatography side, we utilized 50 µm internal diameter columns at an effective flow rate of 125 nl/min during peptide elution, essentially aiming to minimize flow rate while maintaining an adequate column backpressure. It has previously been established that lower flow rates can increase identification rates (Kawashima et al., 2019), due to more efficient ionization and reduced background interference (Zheng et al., 2023).

Balancing depth and throughput is a persistent challenge in SCP, whereas longer gradients can increase the number of proteins identified per cell, such an approach also restricts daily sample throughput. Considering that cell-to-cell variance can be extensive, especially in heterogeneous cell populations, typically dozens to hundreds of single cells need to be acquired in order to be able to draw statistically significant conclusions. In this work, we optimized an analytical gradient using the Vanquish Neo UHPLC which conditions, loads, and initiates the gradient at high pressure (>1,000 bar), before gradually reducing flow rate and pressure to a low level for peptide elution. This concomitantly increases effective peptide elution time across the gradient to ∼90%, while still gaining full benefit from efficient ionization. A caveat of using a setup with sufficiently high column backpressure during low flow rates is that loading of the sample onto the analytical column is relatively time-consuming, which makes it impractical to inject volumes larger than a few microliters. In the HPLC context, using a pre-column setup (Zheng et al., 2023), or potentially a dual-column configuration, could minimize overhead. Another approach to increase sample throughput is to label samples and perform multiplexed analysis (Petrosius et al., 2023; Specht et al., 2021), however, multiplexing strategies are typically expensive, laborious, and subject to ratio compression depending on how conditions and channels are combined within the same multiplex (Hogrebe et al., 2018). Overall, we acquired our SCP data at 40 or 60 SPD, which is on par with contemporary methodology (Bubis et al., 2025; Ye et al., 2025), and sufficient for most standard SCP experiments.

We directly compared the Orbitrap Astral MS with the Orbitrap Astral Zoom MS for analysis of low input samples, and find that the Astral Zoom’s low input mode, pre-accumulation features, and enhanced spectral processing conferred measurable increases in precursor and protein identifications (Fig. 2B). These empirical gains align with what has been observed when using the Astral Zoom in the context of high-load and high-throughput proteomics (Guzman et al., 2025), highlighting that the instrument provides benefits across all proteomics applications.

In the context of SCP or low-input proteomics, some studies rely on FAIMS (Bubis et al., 2025; Petrosius et al., 2023), whereas others acquire data without FAIMS (Mund et al., 2022; Ye et al., 2025). In this study, we designed SCP methodology that works both without and with FAIMS, and achieved similar results from 100-250 pg HeLa lysate, as well as from single HeLa cells. To achieve this without FAIMS, we utilized ultra-sensitive nDIA wherein the precursor scan range is restricted to a width of 200 m/z, allowing the mass spectrometer to spend more time accumulating for each DIA window while keeping the cycle time constant at ∼1 sec. While this comes at a cost of peptides identified, we find that the number of proteins identified was increased, and moreover the quantitative precision was improved for ultra-sensitive nDIA. Further improvements could potentially be made through exclusion of known polysiloxane peaks from the DIA windows, or through overlapping DIA windows to further increase the number of data points per peak.

Application of FAIMS provided a clear advantage at ultra-low loads (<50 pg) by removing singly charged background and enriching for multiply charged peptide ions, yielding up to 2-fold increases in protein identifications from HeLa lysate in the low picogram range, and up to 40% more proteins from single PBMCs. FAIMS has previously been shown to increase protein identifications at high input amounts, albeit at a cost of peptide identifications (Hebert et al., 2018). In the context of low input proteomics, others have also observed that FAIMS increases identifications (Petrosius et al., 2023). Although FAIMS efficiently removes singly-charged contamination, it does come at a modest loss of overall transmission, and we speculate that the cross-over point where acquisition with FAIMS becomes superior likely correlates with both the level of atmospheric contamination within the lab as well as the amount and purity of the samples analyzed. In this context, ongoing efforts towards increasing the transmission of FAIMS could likely expand the range of samples and applications where it would be beneficial to utilize FAIMS (Hoch et al., 2025).

Compared to contemporary SCP studies investigating HeLa or comparably sized cells (Bubis et al., 2025; Ye et al., 2025), we identified significantly more peptides and proteins. We speculate this is due to a combinatory effect of enhanced sample preparation, optimized chromatography, augmented MS acquisition strategy, and utilization of the Orbitrap Astral Zoom MS. Notably, we achieved the highest number of proteins identified at a throughput of 40 SPD, with other labs utilizing 50-80 SPD, or even 120 SPD while retaining solid performance (Ye et al., 2025).

Applying our SCP pipeline to PBMCs, we were able to quantify up to 4,000 proteins per single PBMC (7 – 13 µm), observed strong enrichment for immune and infectious disease pathways, and detected a panel of CD markers sufficient to segregate monocytes and activated lymphocytes by unsupervised clustering. Notably, we did not use computational or experimental carriers to increase identification numbers. Strikingly, our single PBMC proteomic results rivaled large-scale bulk PBMC studies (Giammaria et al., 2025; Koncarevic et al., 2014; Li et al., 2022; Rieckmann et al., 2017; Santa et al., 2024; Shen et al., 2019), while requiring seven orders of magnitude less input material. Of note, the bulk studies had a range of different biomedical aims and relied on a range of different hardware, making it difficult to directly compare. Further, the comprehensive study by Rieckmann et al. identified considerably more proteins overall (Rieckmann et al., 2017), but used approximately three weeks of MS acquisition time. Importantly, as we analyzed single PBMCs, our capability to resolve immune cell identity and pathway coverage provides a direct, label-free molecular alternative to antibody-based cytometry and imaging approaches, and agrees with the notion that proteomics can now directly tackle clinically relevant primary samples.

Taken together, iterative refinement of LC-MS parameters, incorporation of FAIMS, and implementation of the Orbitrap Astral Zoom mass spectrometer enabled ultra-deep profiling of HeLa cells surpassing 7,000 proteins per cell. Furthermore, we achieved comprehensive proteome coverage of PBMCs, identifying ∼4,000 proteins per single PBMC and revealing cell-type-specific signatures directly from mixed populations. These results demonstrate that a carefully optimized SCP workflow can achieve both exceptional depth and throughput without the need for carrier proteomes or labeling strategies. Collectively, this work establishes a robust, accessible platform for single-cell proteomics that bridges the gap between discovery-scale analysis and clinically relevant applications, opening new avenues for exploring cellular heterogeneity and molecular mechanisms in health and disease.

## METHODS

### Cell lines and cell culture

HeLa cells (CCL-2, female) and HEK 293T cells (CRL-3216, female) were acquired via the American Type Culture Collection. All cells were cultured at 37 °C and 5% CO2 in Dulbecco’s Modified Eagle’s Medium (Invitrogen) supplemented with 10% fetal bovine serum and a penicillin/streptomycin mixture (100 U/mL; Gibco). All cells were routinely tested for mycoplasma. Cells were not routinely authenticated.

### Peripheral blood mononuclear cells preparation

Fresh buffy coats were derived from human peripheral blood from anonymous donors, and were acquired from Rigshospitalet, Copenhagen. While these buffy coats were partially enriched for leukocytes, they still contained some red cells and other blood components. Peripheral blood mononuclear cells (PBMCs) were isolated from the buffy coats using Lymphoprep™ (STEMCELL Technologies) density gradient medium, essentially according to the manufacturer’s instructions. Briefly, 15 mL Lymphoprep was added to Leucosep 50 mL tubes (Greiner), and centrifuged for 1 min at 1,000*g* at room temperature. Buffy coats were diluted 1.25x using phosphate buffered saline, and approximately 25 mL of diluted buffy coat was poured into the Leucosep tubes (up to the 40 mL line). The tubes were centrifuged at 1,000*g* for 10 min at room temperature, in a swing-out centrifuge with delayed acceleration and deceleration. Mononuclear cells were harvested by pouring the liquid standing on top of the filter into clean 50 mL polypropylene tubes, after which PBS was added to fill up the tubes. Cells were pelleted via centrifugation at 500*g* for 5 min at room temperature, and the pellet was washed once more in the same manner. Purified PBMCs were resuspended in a mixture of 90% fetal bovine serum and 10% DMSO, prior to being slowly frozen to −80 °C and stored for up to several months until further handling. To recover PBMCs, they were slowly thawed out on ice, after which they were washed once with PBS and pelleted at 1,000*g* to recover a mixed population of PBMCs. Alternatively, they were washed several times with PBS and centrifuged at 500*g*, removing the larger cells that had pelleted, and keeping the supernatant containing only the smallest PBMCs.

### Bulk cell lysis and protein digestion

Cells were washed twice with ice-cold PBS, and gently scraped at 4 °C in a minimal volume of PBS. Cells were pelleted by centrifugation at 500*g*, and vigorously lysed in of SDS Lysis Buffer (1% SDS, 100 mM Tris-HCl, pH 8.5). Lysates were heated to 99 °C for 10 min while shaking vigorously (1,400 RPM) in a ThermoMixer, after which lysates were cooled to 30 °C and centrifuged for 2 min at 14,000*g*. Reduction and alkylation of protein disulfide bonds was performed via concomitant addition of tris(2-carboxyethyl)phosphine (TCEP) and 2-chloroacetamide (CAA) to final concentrations of 10 mM, and incubation at 30 °C for 30 min. On-bead proteolytic digestion was performed using the protein aggregation capture (PAC) protocol, essentially as described previously (Batth et al., 2019). Briefly, proteins were aggregated from the SDS lysate via addition of 10 volumes of acetonitrile (ACN). Magnetic beads (MagReSyn® Hydroxyl) were added to the mixture in a 1:2 protein-to-bead ratio, and samples were mixed three times by gentle inversion, waiting for 1 min in between each mixing. Beads were allowed to settle by gravity for 2 min, prior to being immobilized via a magnetic rack. Immobilized beads were washed twice with 100% ACN, and once with 70% ethanol, after which all liquid was removed and beads were allowed to air-dry for 10 min. Beads were resuspended in 50 mM HEPES pH 8.5, and proteins were digested on-beads via addition of Lys-C (Wako) at a 1:200 enzyme-to-protein (w/w) ratio for 1 h at 37 °C, and subsequently via addition of modified sequencing-grade trypsin (Promega) at a 1:100 (w/w) ratio, overnight at 37 °C. Following digestion, beads were immobilized on the magnet, and digests were clarified through 0.45 µm spin filters prior to peptide cleanup using C18 StageTip.

### StageTip purification of peptides

Bulk HeLa digest peptides were purified using C18 StageTips at low pH. To this end, C18 StageTips were prepared in-house, by layering four plugs of C18 material (Sigma-Aldrich, Empore SPE Disks, C18, 47 mm) per StageTip. Activation of StageTips was performed with 100 μL 100% methanol, followed by equilibration using 100 μL 80% ACN in 0.1% formic acid (FA), and two washes with 100 μL 0.1% FA. Samples were acidified via addition of TFA to a final concentration of 0.5%. Samples were centrifuged for 5 min at 14,000g, after which supernatants were loaded on StageTips. Subsequently, StageTips were washed twice using 100 μL 0.1% FA, after which peptides were eluted using 80 µL 30% ACN in 0.1% formic acid. All samples were dried to completion using a SpeedVac at 60 °C, dissolved to a final concentration of 25 ng/µL in 0.1% FA, and stored at −20 °C. For LC-MS analysis, peptides were diluted 200-fold in 0.1% FA to 125 pg/µL, optionally containing n-dodecyl β-D-maltoside (DDM). For the majority of experiments, DDM was added to a final concentration of 0.005% (50 ng/µL). HeLa experiments were performed using 125 pg (1 µL) or 250 pg (2 µL), and dilution experiments were done via serial dilution ensuring that either 0.5, 1, or 2 µL were injected, in all cases resulting in injection of 25 to 100 ng of DDM per run.

### Single cell isolation and digestion

Cultured cells were washed twice with PBS, and detached from the dish using Trypsin-EDTA (Gibco), which was quenched with a small volume of cell culture medium. Cells were pelleted by centrifugation in a swing-out centrifuge at 500*g* for 2 min, and washed twice with PBS. Cells were diluted to ∼200 cells/μL using PBS, and kept on ice until single-cell sorting. PBMCs were prepared as outlined above, and also diluted to ∼200 cells/µL using PBS and kept on ice until sorting. A CellenOne (Cellenion) was used to dispense ∼500 nL of ice-cold master mix into each well of LoBind 96-well plates (Eppendorf), using a PDC M Piezo Dispensing Capillary (Cellenion) which has an average drop volume of ∼375 pL. Master mix contained 30 mM triethylammonium bicarbonate (TEAB) pH 8.5, 0.05% n-dodecyl β-D-maltoside (DDM), 10 ng/µL Trypsin Gold (Promega; Mass Spectrometry Grade), 5 ng/µL Lys-C (Wako), and 250 µM dithiothreitol (DTT; Proteomics Grade). This master mix was used for the majority of the experiments; any deviations are indicated in figure legends. Mastermix was always kept on ice, and mixed well and centrifuged for 2 min at 14,000*g* just prior to drawing and dispensing. After dispensing of mastermix, the PDC was flushed twice, and used to draw cells. The CellenOne was used to characterize properties of the cell populations, and used to dispense single HeLa (25 – 35 µm), single HEK (20 – 35 µm), or single PBMC (6 – 15 µm) cells directly into the 96-well plate containing mastermix. For most experiments, several wells were configured to receive 20 cells to serve as a ‘carrier’. For specific experiments, either 2, 5, 40, or 60 cells, were pooled into some wells, which is indicated in the figure legends. After sorting, another 250 nL of mastermix was dispensed into all wells. Subsequently, digestion was performed using the CellenOne, at 50 °C for 2 hours, during which the PDC continuously dispensed small volumes of distilled water into each of the wells to both cause a mixing effect within the wells and offset the evaporation occurring at 50 °C. After digestion, the CellenOne was used to cool down the plate to 20 °C for 20 min, during which dispensing of distilled water continued. At all stages of CellenOne sample preparation and depending on atmospheric conditions, the system automatically controlled relative air humidity to prevent either drying out of the wells, or condensation of liquid within the system. This ensured a final volume of ∼4 µL in each well, and following the digestion procedure the plates were briefly centrifuged at 250*g* to ensure all samples were similarly located at the bottom of the wells. Reactions were quenched and stabilized via addition of 1 µL of 1.5% formic acid, after which plates were stored at 4 °C for up to two weeks until MS analysis. Just prior to transferring the plates to the HPLC, plates were centrifuged for 2 min at 250*g*. In the HPLC, plates were kept at 7 °C.

### Liquid chromatography setup

The vast majority of samples were analyzed on 20-cm long analytical LC columns with an internal diameter of 50 µm, packed in-house using ReproSil-Pur 120 C18-AQ 1.9 µm beads (Dr. Maisch), with other column setups defined in Supplementary Table X. On-line reversed-phase liquid chromatography to separate peptides was performed using a Vanquish™ Neo UHPLC System (Thermo Scientific), and the analytical LC column was heated to 40 °C using a column oven (Sonation). Peptides were eluted from the column using a gradient of Buffer A (0.1% formic acid) and Buffer B (80% ACN in 0.1% FA). As optimization of analytical gradient is a main focus in this study, a large number of gradients were used and are described in Supplementary Table X. Expected sample throughput per day (SPD) is indicated in figure legends. Pre- and post-run column equilibration was set to 0.5 column volumes, additional volume of buffer following sample load was set to 1 µL, and all loading and equilibration was performed at 1,200 bar. For the majority of the 60SPD experiments, the gradient was started at 500 nL/min (∼1,000 bar) with 10 %B. Linearly over 1 min, flow rate was reduced to 125 nL/min while increasing to 23 %B, resulting in a stabilized pressure of ∼250 bar after 2 min and synchronized with the start of peptide elution. Subsequently, over 4.75 min to 31 %B, over 2.25 min to 39 %B, and over 1 min to 47 %B, which signified the end of the recorded gradient due to a column delay of ∼6 min at 125 nL/min flow. Next, over 4.75 min to 95 %B, flow rate was linearly increased to 500 nL/min over 0.25 min, and kept there for 1 min. The last ∼6 min of the LC gradient volume eventually passed through and washed the column, but were not recorded by the MS.

### Mass spectrometric analysis

All samples were analyzed using an Orbitrap Astral mass spectrometer or an Orbitrap Astral Zoom mass spectrometer prototype. Electrospray ionization (ESI) was achieved using a Thermo Scientific^TM^ Nanospray Flex^TM^ Ion Source. Spray voltage was set to 2 kV, capillary temperature to 275°C, and RF level to 50%, unless otherwise indicated. As optimization of MS settings is a main focus in this study, a large number of different configurations were used across all experiments, and all the settings are detailed in Supplementary Table X. Here, we describe the settings used for the majority of single-cell proteomics experiments, which were acquired either without (“no-FAIMS”) or with (“FAIMS”) the use of a Thermo Scientific^TM^ FAIMS Pro Duo Interface. All full precursor (MS1) scans were acquired using the Orbitrap™ mass analyzer, while all tandem fragment (MS2 / DIA) scans were acquired in parallel using the Astral™ mass analyzer. Orbitrap lock mass correction was performed at run start using EASY-IC™, for no-FAIMS runs only. For FAIMS, the compensation voltage (CV) was set to −46 V for all scans. Expected peak width was set to 5 s. Full scan range was set to 420-630 m/z (no-FAIMS) or 400-800 m/z (FAIMS), MS1 resolution to 240,000, MS1 AGC target to “500” (5,000,000 charges), and MS1 maximum injection time to 25 ms. DIA scans were executed across a scan range of 425-625 m/z (no-FAIMS) or 400-800 m/z (FAIMS), using 4 m/z (no-FAIMS) or 20 m/z (FAIMS) isolation windows, DIA maximum injection time of 17.5 ms (no-FAIMS) or 40 ms (FAIMS), and fragmentation was performed using HCD with normalized collision energy of 25. DIA fragment scan range was set to 150-1,500 m/z, DIA AGC target to “200” (20,000 charges). Loop control was set to “All”, resulting in a full MS1 & DIA cycle time of ∼1 sec. For acquisition using the Orbitrap Astral Zoom mass spectrometer prototype, pre-accumulation was enabled via the method editor, and higher acquisition rate (HAR), enhanced spectral processing (ESP), and low input mode (LIM) were manually enabled via the Service Diagnostic Tool using a custom license. Low input mode was emulated on the prototype instrument by increasing the PMT voltage by 75 V (∼13%), unless otherwise indicated.

### MS data analysis, general

All MS data files were analyzed with DIA-NN v1.9.2 (Demichev et al., 2020) or Spectronaut 19 (Biognosys). Considering the vast amount of MS data and different experiments, multiple separate computational searches were performed, depending on experimental design. Search engines and search modes are indicated in the figures, figure legends, or otherwise described in Supplementary Table X. Here, we describe the global configuration used for the large majority of all data searches. For generation of theoretical spectral libraries, a complete HUMAN.fasta reference proteome database including isoforms was downloaded from UniProt on the 13^th^ of November, 2024, containing 105,529 entries. Notably, for both DIA-NN and Spectronaut, inference to protein-coding genes was done via gene names, to avoid inflation of identification rates via isoforms.

### MS data analysis, DIA-NN

For analysis with DIA-NN v1.9.2, RAW files were always converted to DIA files in order to speed up subsequent analysis. For library generation for library-free searching, minimum and maximum peptide length were set to 6 and 60 respectively, and minimum and maximum charge to 2 and 4 respectively. Precursor m/z range and fragment m/z range were set to match the MS acquisition strategy, in case of multiple different ranges in the same search, the library range was defined broad enough to encompass all experiments. Missed tryptic cleavages were set to 1 for HeLa lysate searches, or 2 for single cell searches. N-terminal methionine excision was allowed for all searches, and cysteine carbamidomethylation was only used for HeLa lysate searches. For match-between-run library generation, smart profiling was used for HeLa lysate searches and full profiling was used for single cell searches. Quantification strategy was set to “Legacy”. Default settings were used otherwise, including protein-group inference via unique gene name and FDR control set at 1%.

### MS data analysis, Spectronaut

For analysis with Spectronaut 19, default settings were used, with exceptions and notable settings outlined below. For the initial Pulsar search, maximum variable modifications per peptide were set to 3, allowing protein N-terminal acetylation, deamidation of glutamine and asparagine, and methionine oxidation. N-terminal methionine excision was allowed (default). Cysteine carbamidomethylation was not used as a fixed modification, as we did not alkylate single cell samples. Missed tryptic cleavages were set at 2 (default), and minimum and maximum peptide length were set to 6 and 60 respectively. Fragment ion m/z range was set to match the MS acquisition strategy. Relative intensity threshold for fragment ions was disabled for searching including non-FAIMS data. Searches were either performed in “Method evaluation” or standard directDIA mode, as indicated in the figure legends. In general, method evaluation was used when combining vastly different datasets in the same computational search. Protein quantification mode was set to “MaxLFQ”, with quantification performed at the MS2 level unless otherwise specified. Major (protein) grouping was performed by gene name (Gene Id), and minor (peptide) grouping was performed by stripped sequence. An FDR of 1% was applied at the PSM, peptide, and protein levels.

### Comparison to other MS studies

For comparison of proteins identified from single HeLa, HEK293T, and H460 cells, publicly available MS data was downloaded from two recent single cell proteomics studies (Bubis et al., 2025; Ye et al., 2025). Specifically, HeLa single cell data was acquired from Ye et al., and HEK293T and H460 data was acquired from Bubis et al. For comparison using Spectronaut, each dataset from each study (i.e. each separate data column in the figure) was set as a separate condition for Method Evaluation analysis using otherwise the same settings as described above. For comparison using DIA-NN, datasets from each study were searched separately using MBR mode, and afterwards combined into one graph. Peptides per protein and datapoints per peak were derived from Spectronaut analysis. For depth of sequencing analysis, protein copy-numbers (via iBAQ) were derived from a previously published ultra-deep proteome, and proteins identified from single cells were aligned to the deep proteome primarily through Uniprot ID, and otherwise through gene name. For comparison of proteins identified from single PBMCs in this study to published bulk PBMC studies, reported data was taken from six other studies (Giammaria et al., 2025; Koncarevic et al., 2014; Li et al., 2022; Rieckmann et al., 2017; Santa et al., 2024; Shen et al., 2019). In order to align all identified and reported proteins, a scaffold was created based on all unique protein-coding genes as downloaded from UniProt on the 13^th^ of November, 2024, i.e. the same database as used to search all MS data in this work. Alignment was primarily performed using Uniprot identifier, and otherwise protein-coding gene. Duplicate proteins, or proteins no longer existing on Uniprot, were omitted. For Končarević et al., all reported Uniprot identifiers were aligned. For Rieckmann et al., the vast majority of gene names were aligned, omitting only entries not quantifiable in any of the 43 different conditions, as well as duplicate entries. For Shen et al., all quantified proteins were aligned. For Li et al., since only unfiltered MaxQuant output was available, we performed basic filtering by removing contaminants and reverse database hits, filtering for at least 2 peptides per protein, proteins identified in less than 60% of replicates were discarded; as also performed by Li et al. For Santa et al., all reported Uniprot identifiers were aligned. For Giammaria et al., since we could not find a list of proteins, we downloaded the publicly available MS raw data associated with their work, and re-analyzed the data using DIA-NN v1.9.2 using the same settings as used for data generated in this study. From this, we found essentially the same number of proteins as the authors reported, and those proteins were aligned to the scaffold for comparison between studies. Our own single PBMC data was searched with both DIA-NN and Spectronaut, and for the study comparison we only used protein-coding genes identified by both software. For determination of which markers occur in which types of PBMC, we used the in-depth quantitative data from Rieckmann et al.

### Quantification and statistical analysis

Details regarding the statistical analysis can be found in the respective figure legends. All quantitative experiments were performed using at least four replicates to ensure sufficient statistical power. For single cell analysis, we aimed to analyze at least 10 single cell replicates for each condition to account for cell-to-cell biological variance in asynchronous cell populations. For some conditions, due to technical issues, we could not measure at least 10 single cells, in which case we drew no statistical conclusions from those datasets. Statistical handling of the data was primarily performed using the freely available Perseus software (Tyanova et al., 2016), and includes calculation of Pearson correlation and linear regression, hierarchical clustering, and dimensionality reduction (UMAP) analysis. Venn diagram overlaps were performed using Venny (https://bioinfogp.cnb.csic.es/tools/venny/), and scaled Venn diagrams were rendered using eulerAPE (https://www.eulerdiagrams.com/eulerAPE/v2/). UpSet plots were generated using ChiPlot (https://www.chiplot.online/). Boxplots were drawn using BoxPlotR (http://shiny.chemgrid.org/boxplotr/). Violin plots were drawn using GraphPad Prism. Term enrichment analyses (Uniprot keywords, Reactome, TISSUES) were performed using the STRING database (Szklarczyk et al., 2019). The “T Cell Receptor Signaling Pathway” was adapted from Kyoto Encyclopedia of Genes and Genomes (KEGG) (Kanehisa and Goto, 2000). Reactome Pathway enrichment analysis was performed using and curated by Reactome (Milacic et al., 2024). Overall, plotting of graphs was done using either Perseus, GraphPad Prism, or Microsoft Excel. Some art was acquired through BioRender. The majority of art was drawn by the authors, and all visualizations were polished using Adobe Illustrator.

## ACKNOWLEDGEMENTS

Work at The Novo Nordisk Foundation Center for Protein Research (CPR) is funded in part by a donation from the Novo Nordisk Foundation (NNF14CC0001, NNF24SA0098829 and NNF21OC0072070). This project was supported by a center-of-excellence grant from the Danish National Research Foundation to Copenhagen Center for Glycocalyx Research (DNRF196). This project was also supported by a grant from the Danish Agency of Higher Education and Science to establish the PLATO research infrastructure: Danish National Mass Spectrometry Platform for Proteomics and Biomolecular Imaging (5229-00012B). J.V.O. and M.Y.L. are also funded by Novo Nordisk A/S (CELFFI-2022-002843).

## AUTHOR CONTRIBUTIONS

I.A.H., S.C.B-L. and J.V.O. designed the experiments. I.A.H., S.C.B-L., T.N.A., and M.Y.L. prepared samples and performed proteomics experiments. I.A.H., S.C.B-L., and M.R. analyzed the resulting data. A.H.K. provided consult. I.A.H., S.C.B-L., and J.V.O. wrote the first draft of the manuscript. D.H. and E.D. critically evaluated the results. All authors read, edited and approved the final version of the manuscript.

## SUPPLEMENTARY FIGURE LEGENDS

**Figure S1.**
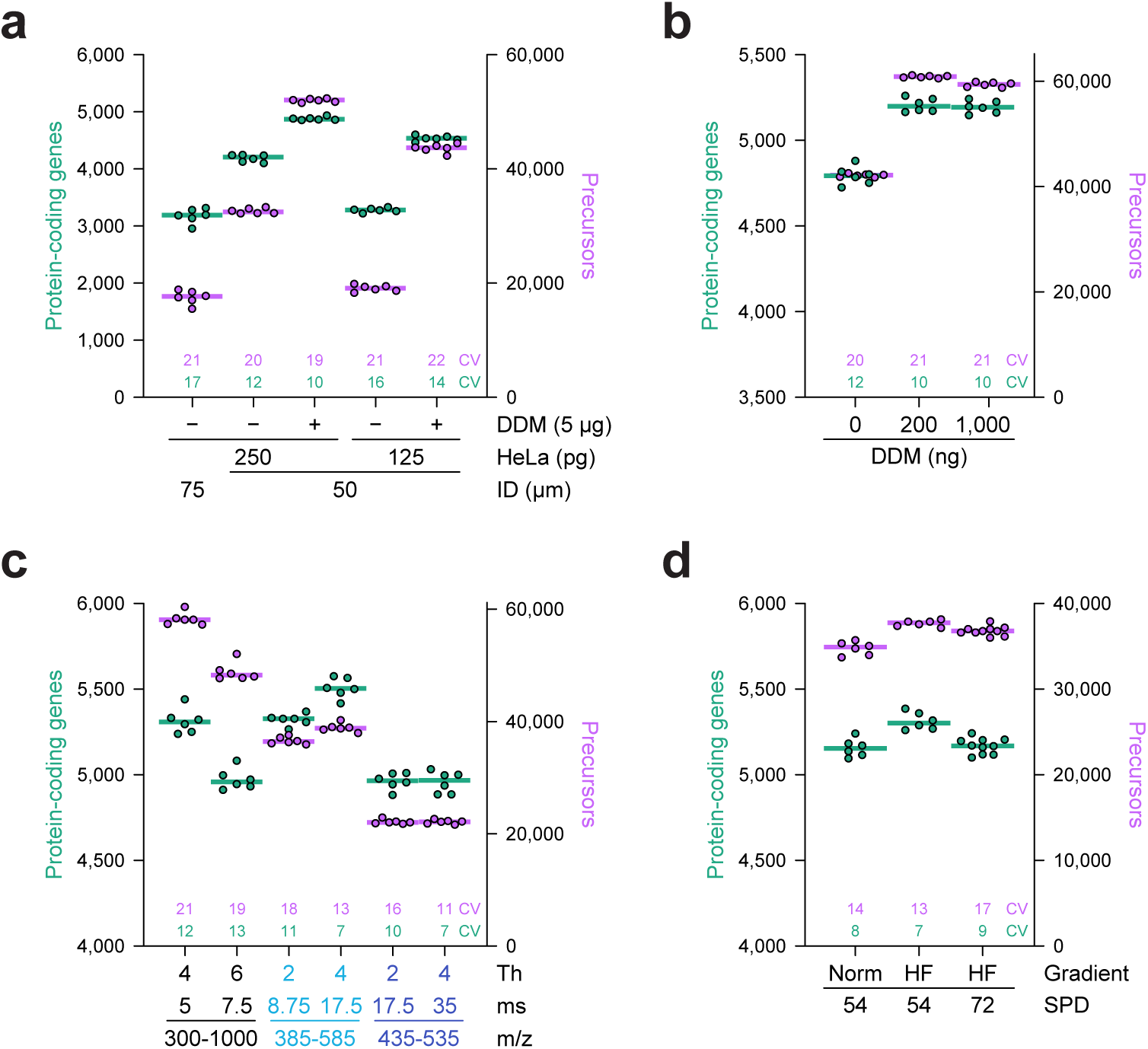
Optimization of low-input LC-MS methodology for enhanced protein identification rates. Corresponds to Figure 1. (A) Number of protein-coding genes (green, left axis) and precursors (purple, right axis) identified, when comparing different column inner diameter, HeLa load, and the addition of n-dodecyl β-D-maltoside (DDM). Bars represent median, the numbers above the x-axis represent median coefficient of variation percentage, *n* = 6 technical replicates. “ID”; internal column diameter. Data analysis using DIA-NN v1.9.2 and matching-between-runs enabled. (B) As **A**, but comparing different amounts of DDM injected. (C) As **A**, but comparing different ultra-sensitive nDIA methods. “ms”; maximum DIA ion injection time in milliseconds. (D) As **A**, but evaluating different gradient lengths and throughput speeds. *n* = 6-10 technical replicates. “Norm”; standard steady-flow chromatography, “HF”; method employing High Flow at the start and end of the gradient to maximize effective peptide elution time, “SPD”; samples per day.

**Figure S2.**
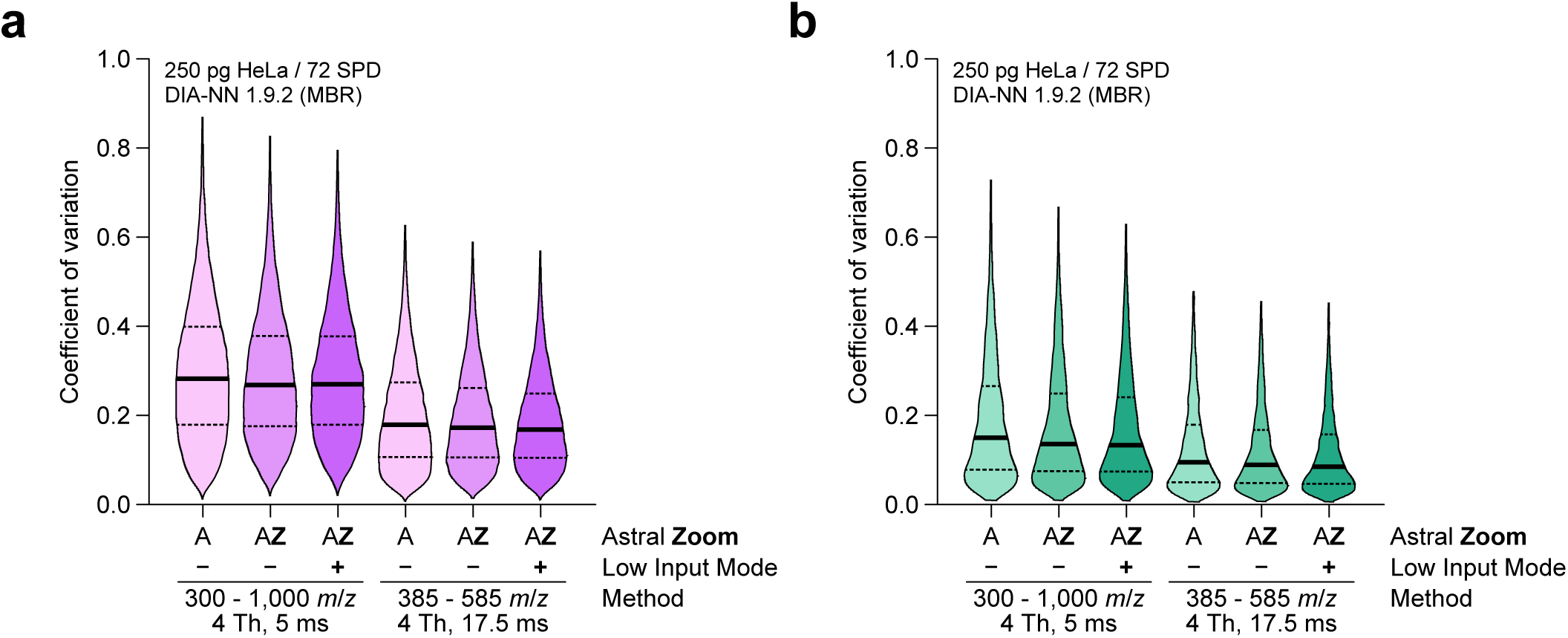
Corresponds to Figure 2B. (A) Violin plot distribution of precursor coefficient of variation observed when comparing “Astral” mode with “Astral Zoom” mode, with the latter either in regular or Low Input mode. Data analysis using DIA-NN v1.9.2 without matching-between-runs, with quantification at the MS/MS level. (B) As **A**, but for protein-coding genes.

**Figure S3.**
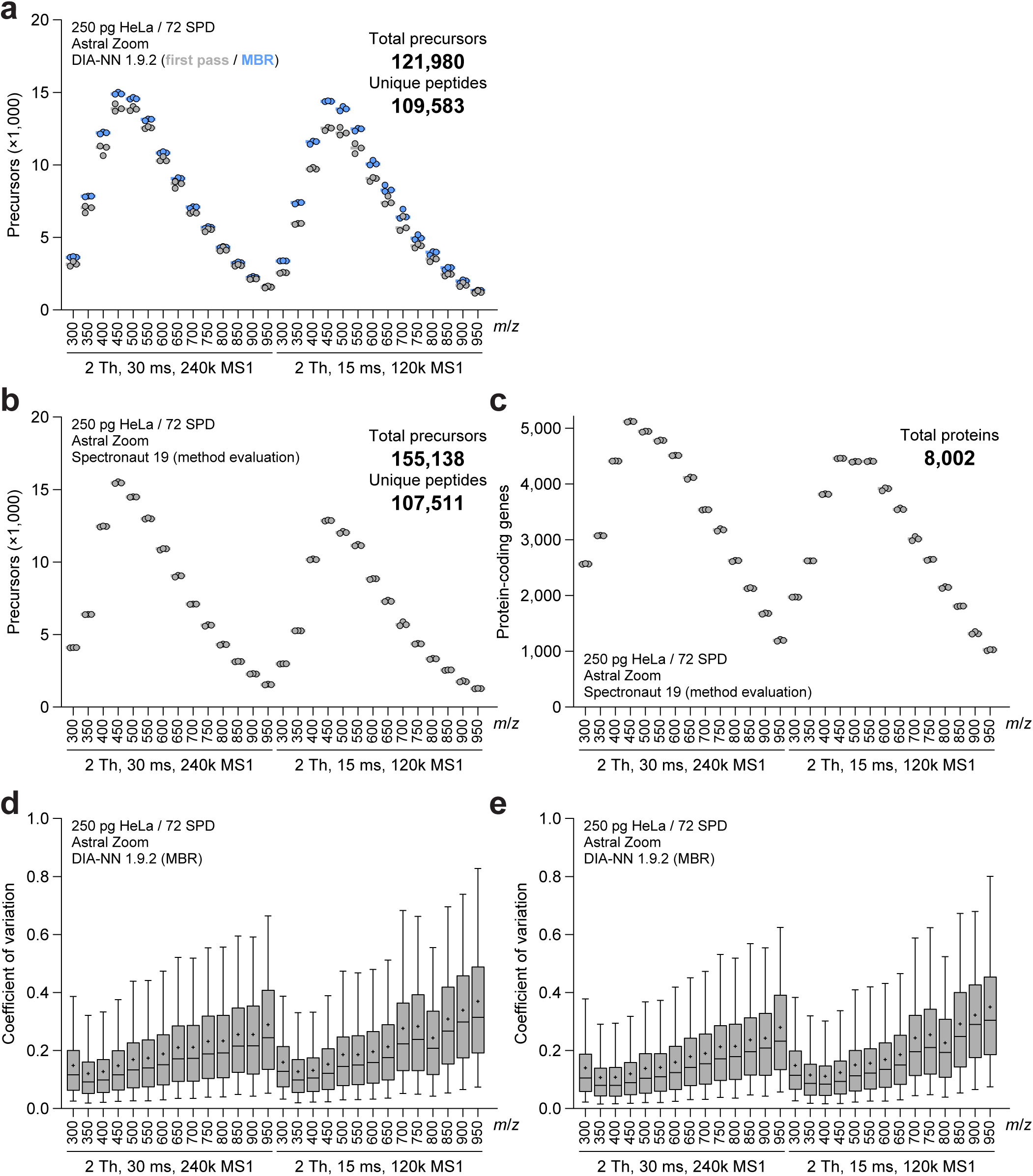
Corresponds to Figure 2C. (A) Gas-phase fractionation experiment, wherein precursors are isolated from a 50 *m/z* range starting at the *m/z* indicated on the x-axis. Gray and blue dots correspond to the number of precursors with data analysis using DIA-NN in first pass mode and MBR mode, respectively. *n* = 3 technical replicates, total number of precursors and unique peptides identified cumulatively across all runs are indicated. (B) As **A**, but data analysis using Spectronaut 19. (C) As **B**, but for protein-coding genes. Total number of protein-coding genes identified cumulatively across all runs is indicated. (D) As **A**, but visualizing coefficient of variation for all identified precursors as a box plot. Central line; median, plus symbol; average, box limits; 1^st^ and 3^rd^ quantile, whisker limits; 5^th^ and 95^th^ percentile. Data analysis using DIA-NN v1.9.2 with matching-between-runs, with quantification at the MS/MS level. (E) As **D**, but for protein-coding gene coefficient of variation.

**Figure S4.**
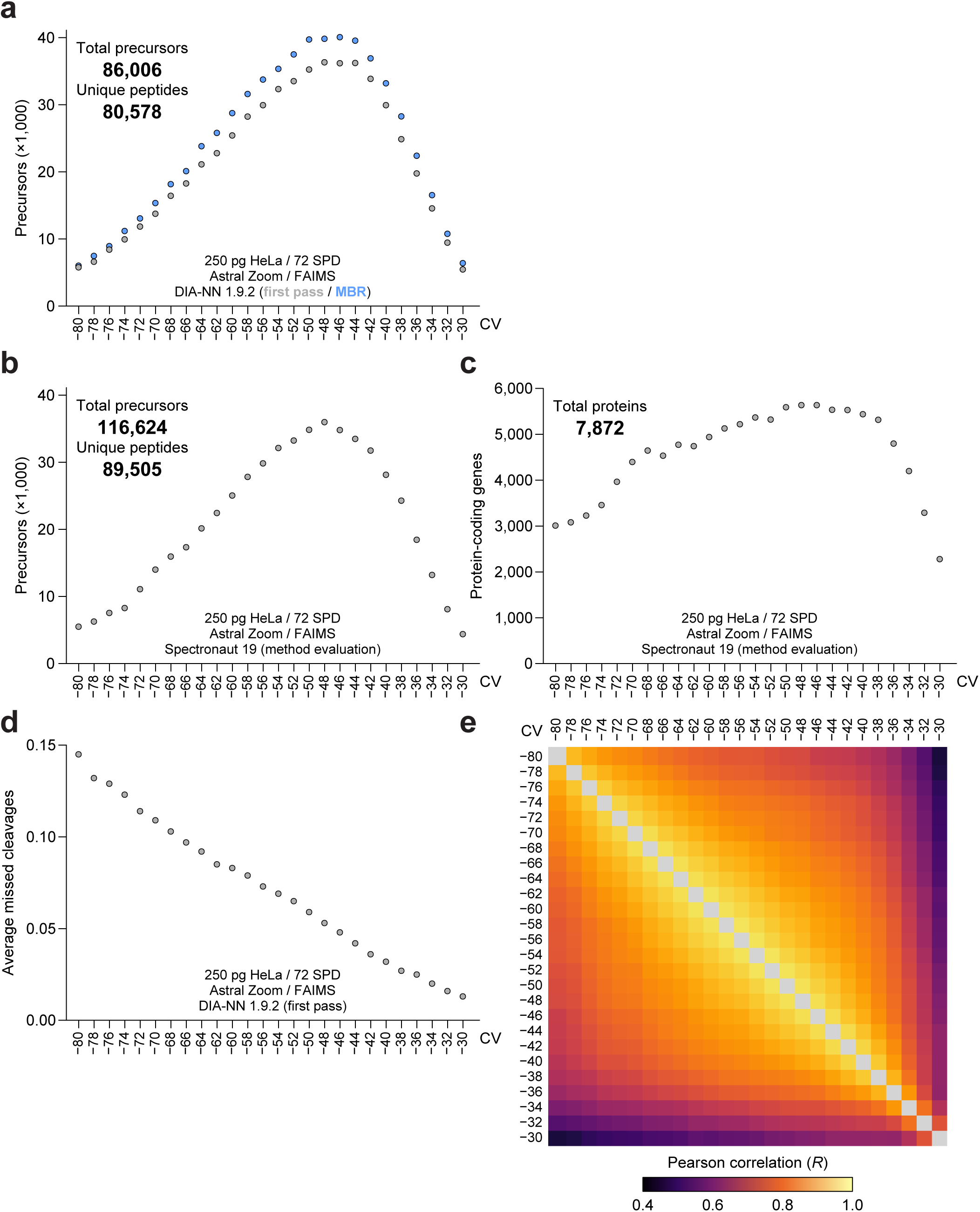
Corresponds to Figure 2D. (A) FAIMS fractionation experiment, wherein precursors were analyzed using a range of FAIMS compensation voltages (CV), using a single constant CV across the entire gradient. Gray and blue dots correspond to the number of identified precursors with data analysis using DIA-NN v1.9.2 in first pass mode and MBR mode, respectively. *n* = 1 technical replicate, total number of precursors and unique peptides identified cumulatively across all runs are indicated. (B) As **A**, but data analysis using Spectronaut 19. (C) As **B**, but for protein-coding genes. Total number of protein-coding genes identified cumulatively across all runs is indicated. (D) As **A**, but visualizing the average number of missed tryptic cleavages in all identified peptide sequences. (E) As **A**, showing the Pearson correlation between the abundance of all protein-coding genes identified at each FAIMS CV setting.

**Figure S5.**
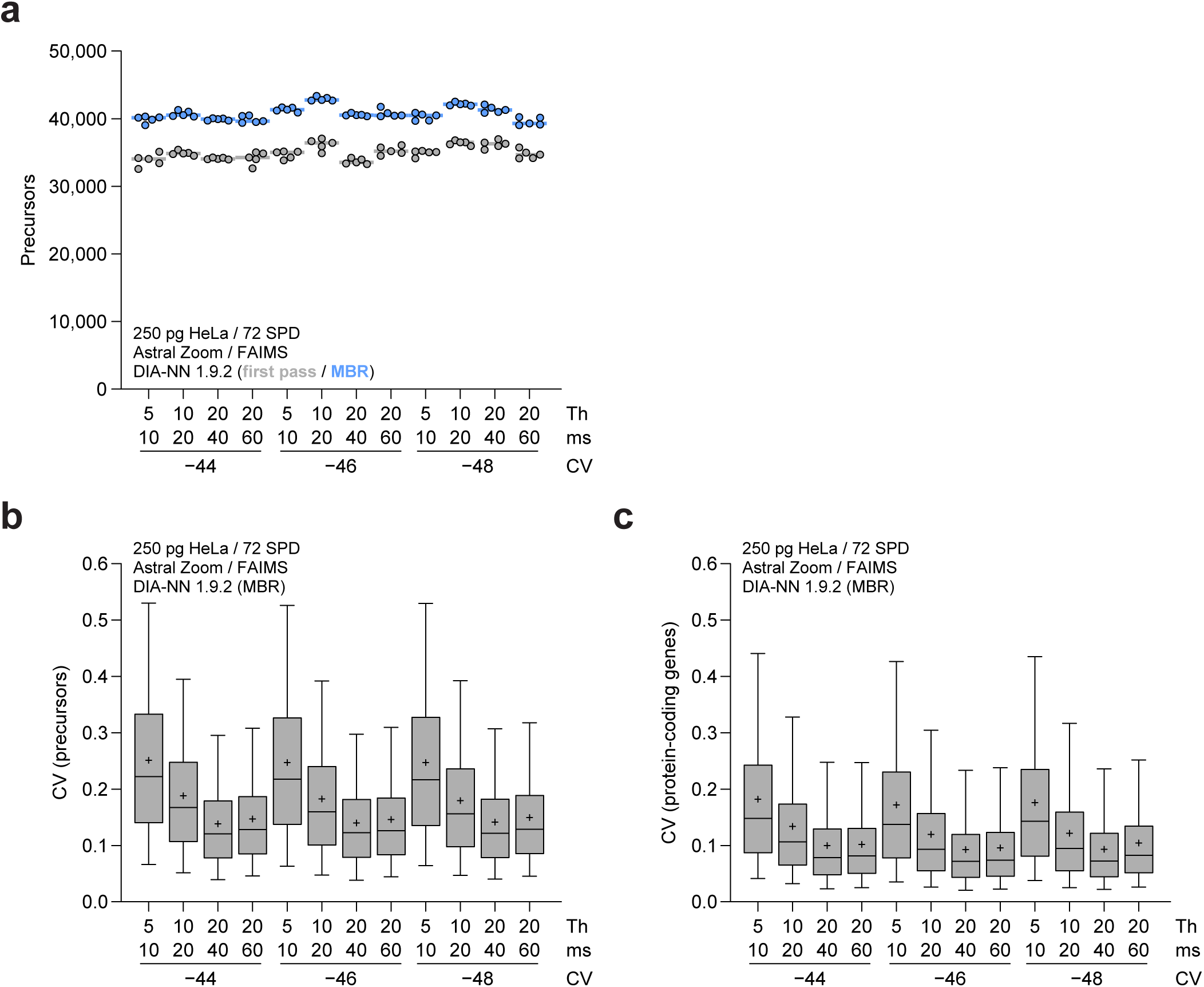
Corresponds to Figure 2E. (A) Number of precursors identified using different combinations of DIA window size in Thomson (“Th”) and maximum allowed DIA ion injection time in milliseconds (“ms”) in combination with three different FAIMS CVs. *n* = 5 technical replicates. Gray and blue dots correspond to data analysis using DIA-NN v1.9.2 in first pass mode and MBR mode, respectively. (B) As **A**, but visualizing coefficient of variation for all identified precursors as a box plot. Central line; median, plus symbol; average, box limits; 1^st^ and 3^rd^ quantile, whisker limits; 5^th^ and 95^th^ percentile. Data analysis using DIA-NN v1.9.2 with matching-between-runs, with quantification at the MS/MS level. (C) As **B**, but for protein-coding gene coefficient of variation.

**Figure S6.**
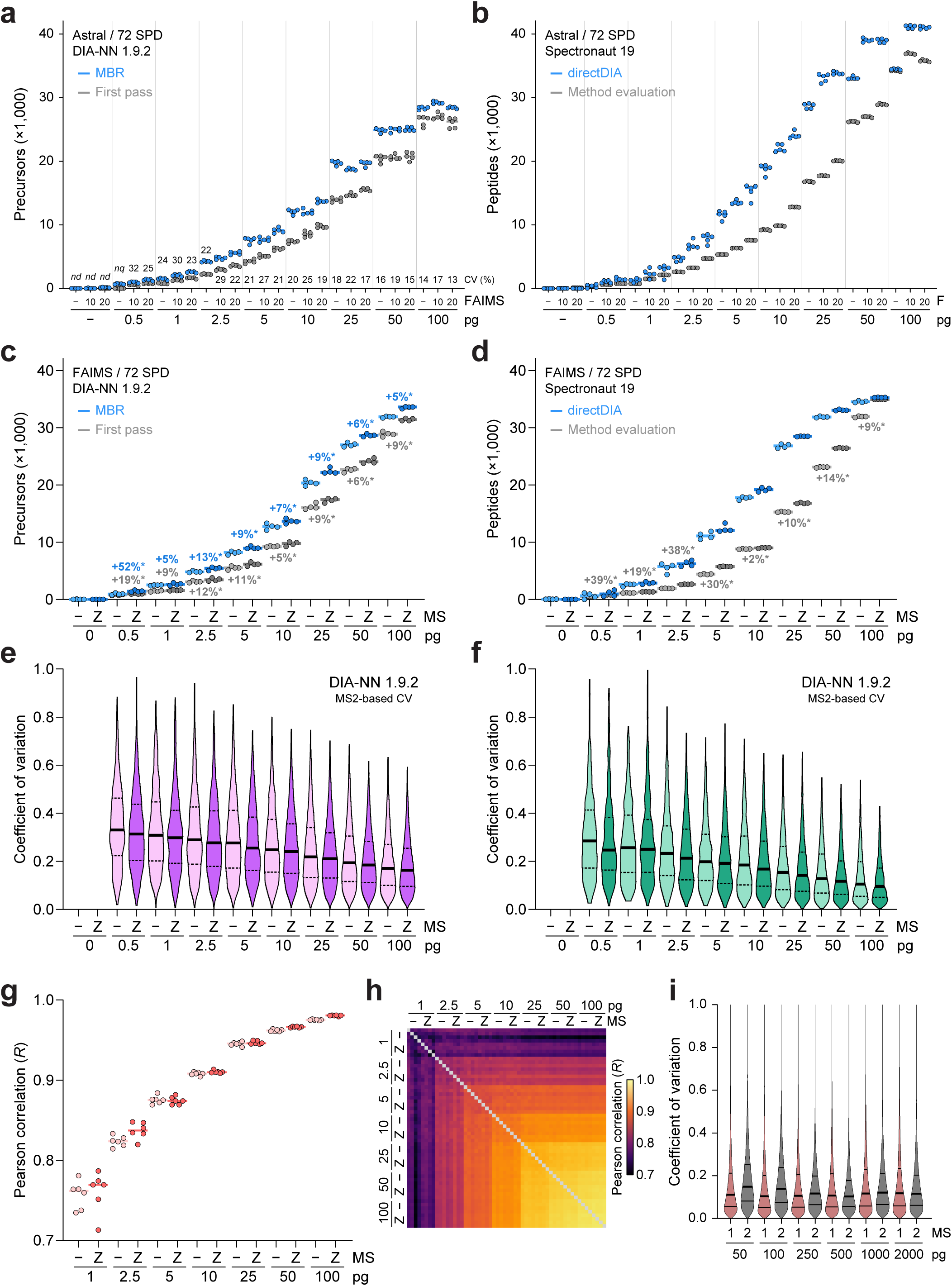
Orbitrap Astral Zoom MS and FAIMS are superior for ultra-low load analysis. Corresponds to Figure 3. (A) Dilution series experiment, ranging from 0.5 pg to 100 pg of injected HeLa lysate, comparing standard analysis to FAIMS analysis. All injections were performed in ascending amounts, starting with a blank to ensure negligible sample carry-over. Number of identified precursors are indicated, bars represent median, with gray and blue corresponding to data analysis using DIA-NN v1.9.2 in first pass mode and MBR mode, respectively. Numbers above the x-axis represent median coefficient of variation percentage, *n* = 5 technical replicates. “F”; indicator for analysis strategy with “−”; no FAIMS, “10”; 10 Th window width and 20 ms DIA injection time, “20”; 20 Th window width and 40 ms DIA injection time. (B) As **A**, but data analysis using Spectronaut 19 and showing number of stripped peptide sequences, with gray and blue corresponding to data analysis using Method Evaluation mode and directDIA mode, respectively. (C) Dilution series experiment as described in **A**, but using FAIMS in all cases and comparing the difference between the Orbitrap Astral and Orbitrap Astral Zoom using a prototype mass spectrometer. Percentage increases of median values are indicated, * = *p* < 0.05 via two-tailed Student’s t-testing, *n* = 4 technical replicates. (D) As **C**, but data analysis using Spectronaut 19 and showing number of stripped peptide sequences, with gray and blue corresponding to data analysis using Method Evaluation mode and directDIA mode, respectively. Differences of medians were not assessed in directDIA mode due to the non-linear behavior of the directDIA identification strategy. (E) Coefficient of variation for all identified precursors in **C**, as a box plot. Central line; median, plus symbol; average, box limits; 1^st^ and 3^rd^ quantile, whisker limits; 5^th^ and 95^th^ percentile. Data analysis using DIA-NN v1.9.2 without matching-between-runs, with quantification at the MS/MS level. (F) As **E**, but for protein-coding genes. (G) Showing the Pearson correlation between the abundance of all protein-coding genes identified in **F** (and Figure 3D), when considering same-replicate correlation only. *n* = 4 technical replicates, leading to 6 combinations (data points), bars indicate the median. (H) As **G**, but comparing the abundance of all protein-coding genes between all replicates. (I) Violin plot distribution of coefficient of variation for all precursors from Figure 3G. MS1-based quantification via Spectronaut, MS2-based quantification via DIA-NN. Thick line represents median, thin lines 1^st^ and 3^rd^ quartiles.

**Figure S7.**
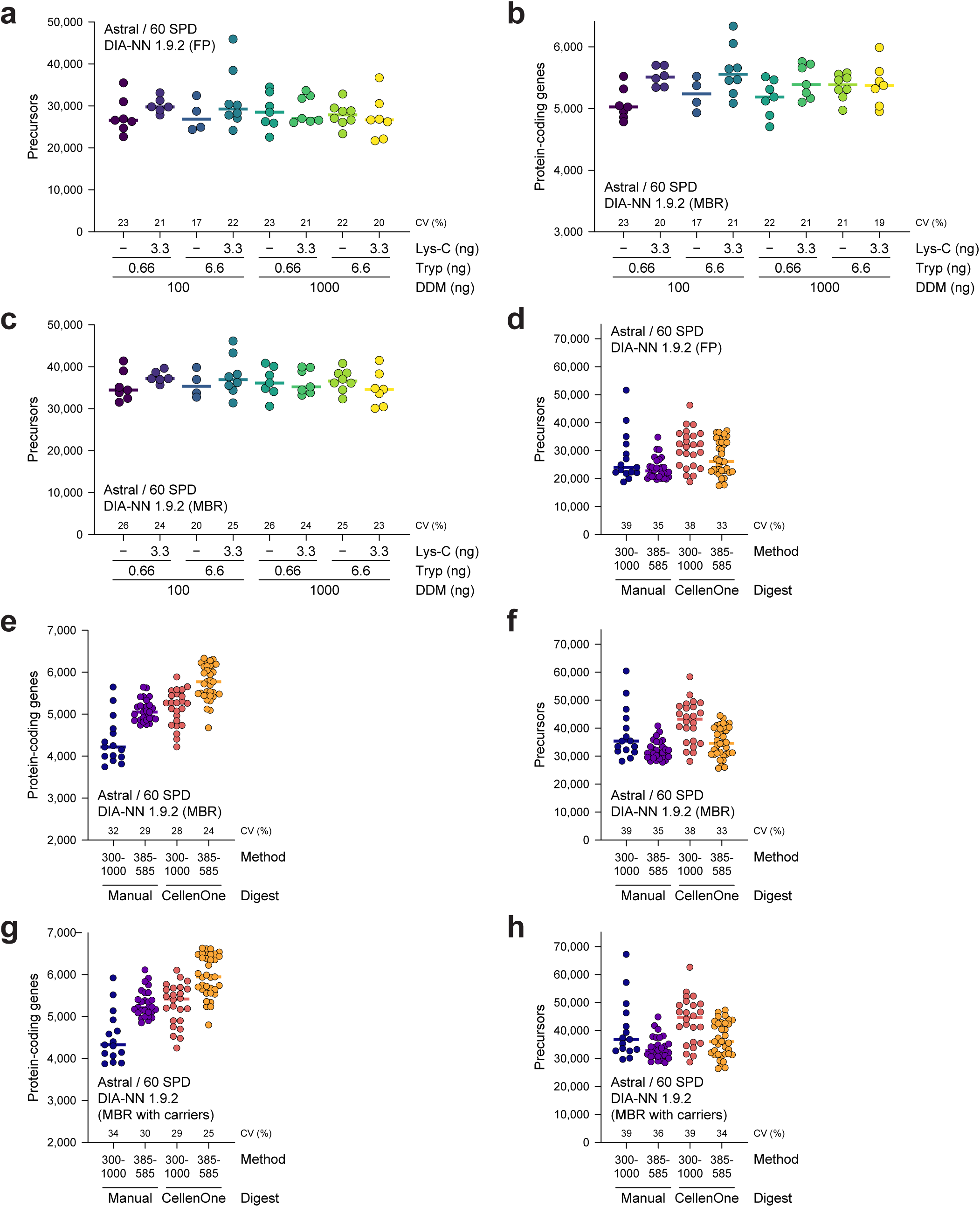
Corresponds to Figure 4B-E. (A) Number of precursors identified from single HeLa cells when using various different digestion mixes. Cells were sorted by the CellenOne platform, but otherwise manually digested. Bars represent median. Numbers above the x-axis are median coefficient of variation. *n* = 4-8 single cells. (B) As **A**, but for protein-coding genes and using matching-between-runs in DIA-NN. (C) As **A**, but using matching-between-runs in DIA-NN. (D) As **A**, but comparing manual digestions (with the CellenOne only used to dispense single cells) to fully automated handling and digestion using the CellenOne, as well as comparing the broad-range nDIA method with ultra-sensitive nDIA. *n* = 15-35 single cells. (E) As **D**, but for protein-coding genes and using matching-between-runs in DIA-NN. (F) As **D**, but using matching-between-runs in DIA-NN. (G) As **E**, but also computationally including 20-cell carriers. (H) As **F**, but also computationally including 20-cell carriers.

**Figure S8.**
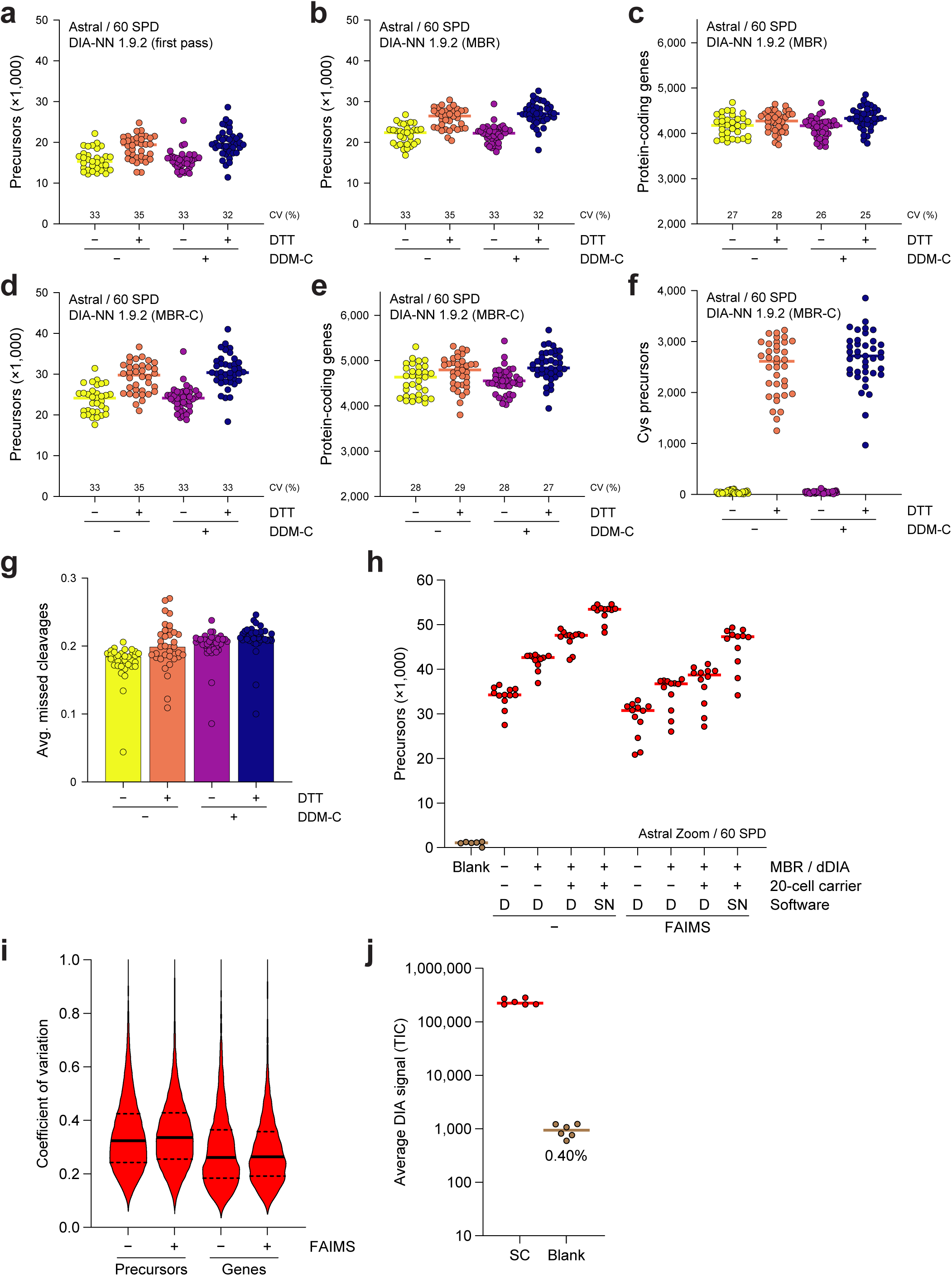
Corresponds to Figure 4F-H. (A) Number of precursors identified from single HeLa cells when comparing the effect of dithiothreitol (DTT) in the digestion mix, as well as pre-coating the 96-well plate with n-dodecyl-ß-D-maltoside (DDM). Cells were sorted by the CellenOne platform, but otherwise manually extracted and digested. Bars represent median. Numbers above the x-axis are median coefficient of variation. *n* = 33-40 single cells. (B) As **A**, but using matching-between-runs in DIA-NN. (C) As **B**, but for protein-coding genes. (D) As **B**, but also computationally including 20-cell carriers (“MBR-C”). (E) As **C**, but also computationally including 20-cell carriers. (F) As **D**, but only showing cysteine-containing precursors. (G) As **A**, but showing the average number of missed tryptic cleavages observed per peptide. (H) Number of precursors (with DIA-NN) or stripped peptide sequences (with Spectronaut) identified from single HeLa cells (median diameter ∼26.5 μm) when using optimized sample preparation workflows, computational inclusion of 20-cell carriers, and using either standard-or FAIMS-optimized methodology. Data analysis using either DIA-NN (“D”) or Spectronaut (“SN”) as indicated. *n* = 12 single cells. “MBR”; DIA-NN matching-between-runs, “dDIA”; Spectronaut directDIA. (I) Violin plot distribution of coefficient of variation for all precursors and protein-coding genes from **H** and Figure 4H. MS2-based quantification via DIA-NN. Thick line represents median, dashed lines 1^st^ and 3^rd^ quartiles. (J) Approximation of system carry-over effect via average DIA ion current signal observed across the entire chromatographic gradient. “SC”; representative single HeLa cell runs. “Blank”; blank injections directly following SC runs with same volume, buffers, and DDM concentration. *n* = 6 single cells and blanks.

**Figure S9.**
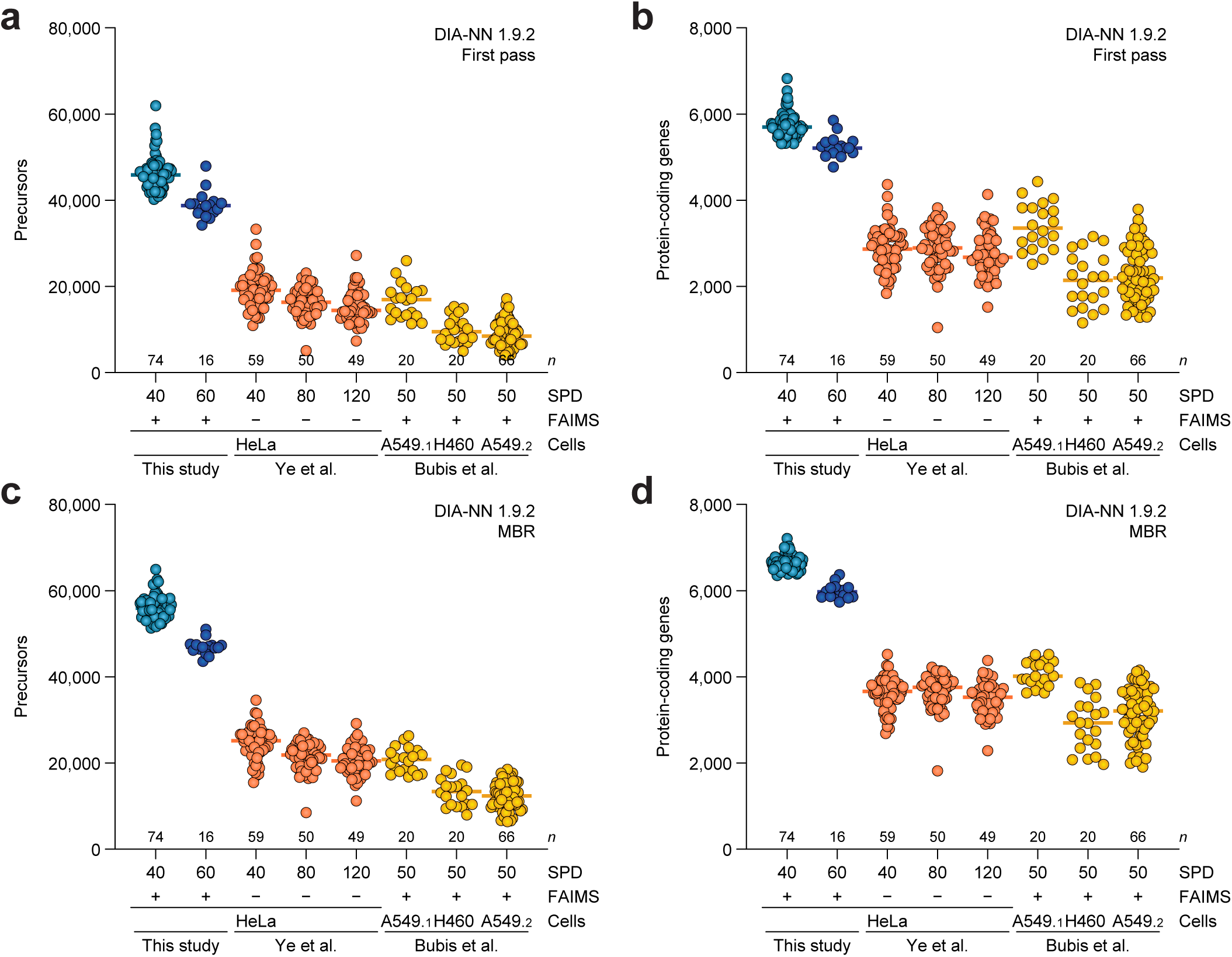
Profiling over 7,000 proteins from single HeLa cells. Corresponds to Figure 5. (A) Comparison between the number of precursors identified from single cells in this study, to two recent papers using similar single-cell approaches (Bubis et al., 2025; Ye et al., 2025). Publicly available MS raw data was downloaded and re-processed using DIA-NN in first pass mode. Bars represent median values. *n* = 16-74 single cells depending on study and batch (see Figure 5B). Appendix numbers following cell type indicate two separate batches of A549 cells from Bubis et al. (B) As **A**, but for protein-coding genes. (C) As **A**, but using matching-between-runs in DIA-NN. To control search space and FDR, each batch of cells was searched separately. (D) As **C**, but for protein-coding genes.

**Figure S10.**
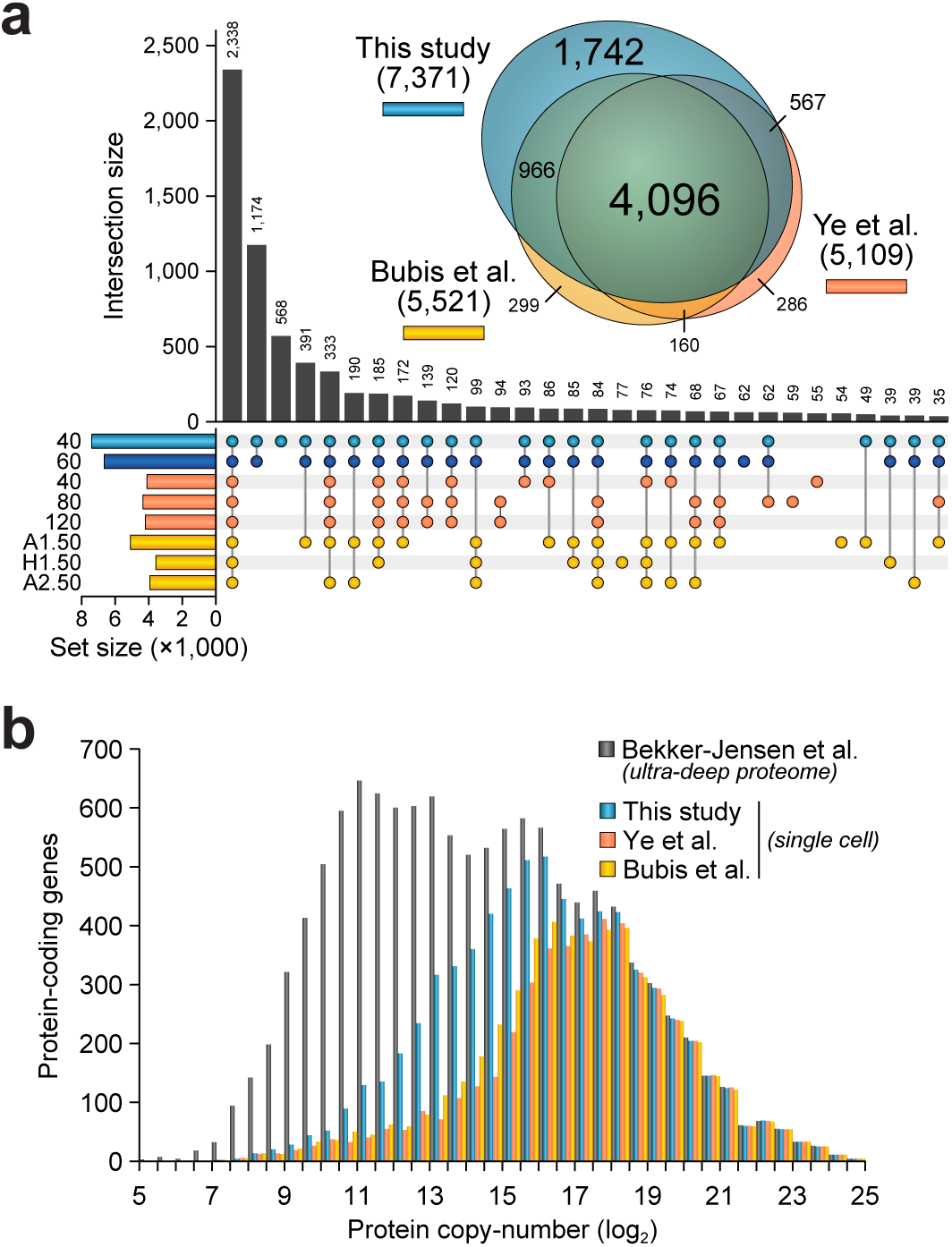
Greatly enhanced single cell proteome depth compared to contemporary single cell studies. Corresponds to Figure 6. (A) UpSet plot analysis, visualizing distinct and overlapping sets of proteins identified from single HeLa cells in this study compared to two recent studies using similar single-cell approaches (Bubis et al., 2025; Ye et al., 2025), see also Figure 5C-F. Exact values are displayed above each bar. Analysis based on re-processing of available MS data using Spectronaut in Method Evaluation mode, considering all proteins (even those identified by just one peptide) detected in at least 50% of replicates. All single cells were HeLa, except for “A1”; A549 batch 1, “H1”; H460 batch 1, “A1”; A549 batch 2. The in-set scaled Venn diagram depicts global overlap between the three studies. (B) Depth of sequencing visualization, with known protein copy-numbers taken from a published ultra-deep proteome. Proteins identified in all single-cell studies were mapped to the total proteome and binned in 0.5 log_2_ bins.

**Figure S11.**
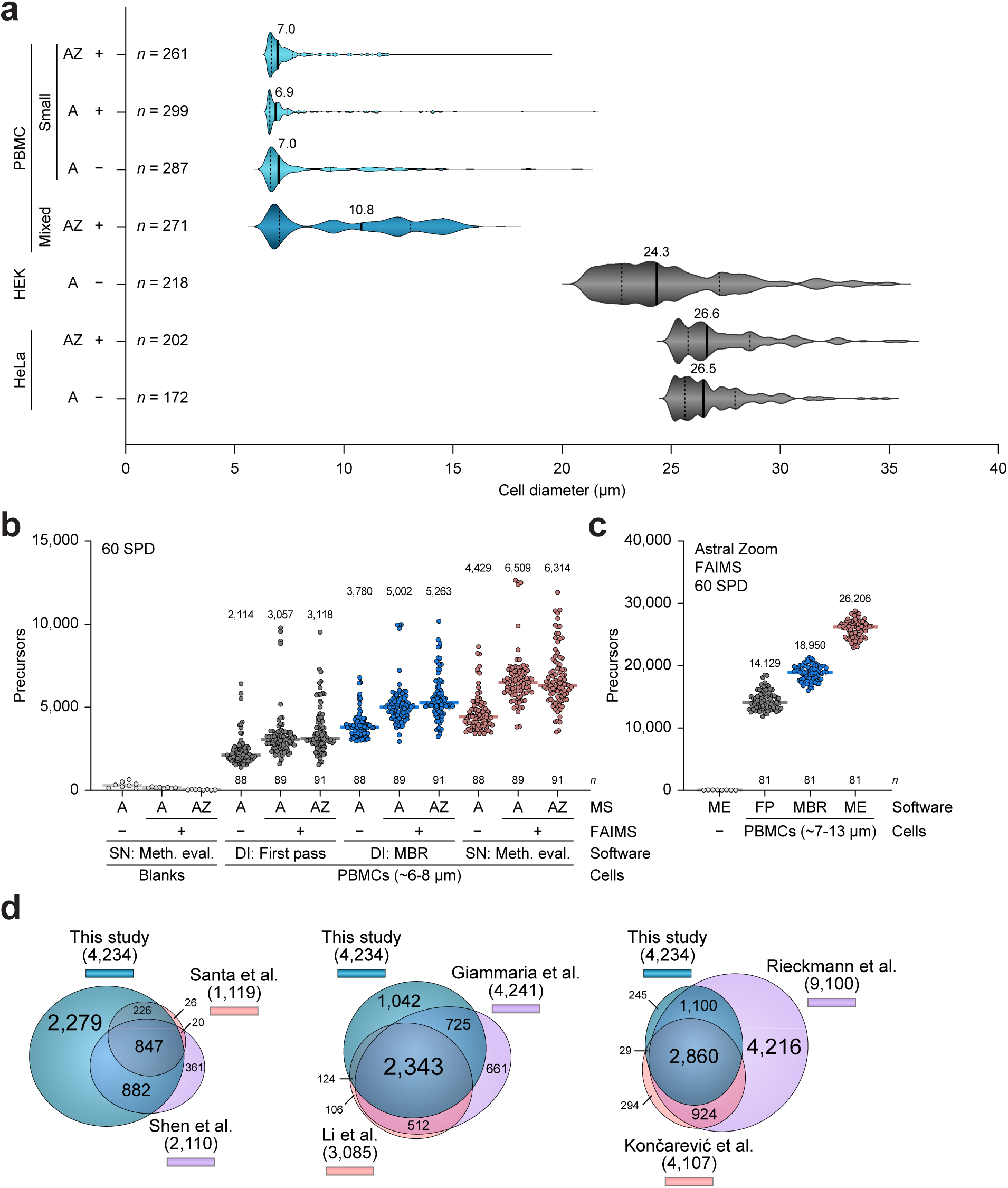
Profiling single peripheral blood mononuclear cells to a bulk analysis depth. Corresponds to Figure 7. (A) Size distribution profile of all PBMCs isolated in this study, stratified by batch used for each set of MS experiments. PBMCs were prepared to exclusively yield the smallest cells (“Small”), or minimally processed to gain a mixed population (“Mixed”). Size distribution profiles of HeLa cells (Figures 4-5) and HEK cells (Supplementary Note **3**) are shown for reference. Thick lines represent median, with median values indicated, and thin dashed lines indicate 1st and 3rd quartiles. *n* = number of sorted cells as indicated. “A”; analyzed by Orbitrap Astral, “AZ”, analyzed by Orbitrap Astral Zoom. A plus symbol denotes analysis using FAIMS. (B) Number of precursors (DIA-NN) or stripped peptide sequences (Spectronaut) identified from the smallest PBMCs (median diameter ∼7 μm), comparing performance of an Orbitrap Astral (“A”) with and without FAIMS, to an Orbitrap Astral Zoom (“AZ”) with FAIMS. Bars represent median values, exact medians displayed above each group of data. Data analysis using DIA-NN (“DI”) or Spectronaut (“SN”) as indicated. “MBR”; DIA-NN matching-between-runs, “dDIA”; Spectronaut directDIA. *n* = 88-91 single PBMCs. (C) As **B**, but visualizing analysis of the Mixed PBMC population using the Orbitrap Astral Zoom with FAIMS. *n* = 81 single PBMCs. “ME”; Spectronaut with Method Evaluation, “FP”; DIA-NN with first pass, “MBR”; DIA-NN with matching-between-runs. (D) Scaled Venn diagrams depicting overlap between proteins identified from single PBMCs in this study, to six bulk PBMC studies. To ensure stringency, proteins identified from single PBMCs in this study were only considered if identified by both DIA-NN and Spectronaut. Proteins from other studies were taken as reported by the respective authors, with data filtering and alignment steps described in the methods section.

**Figure S12.**
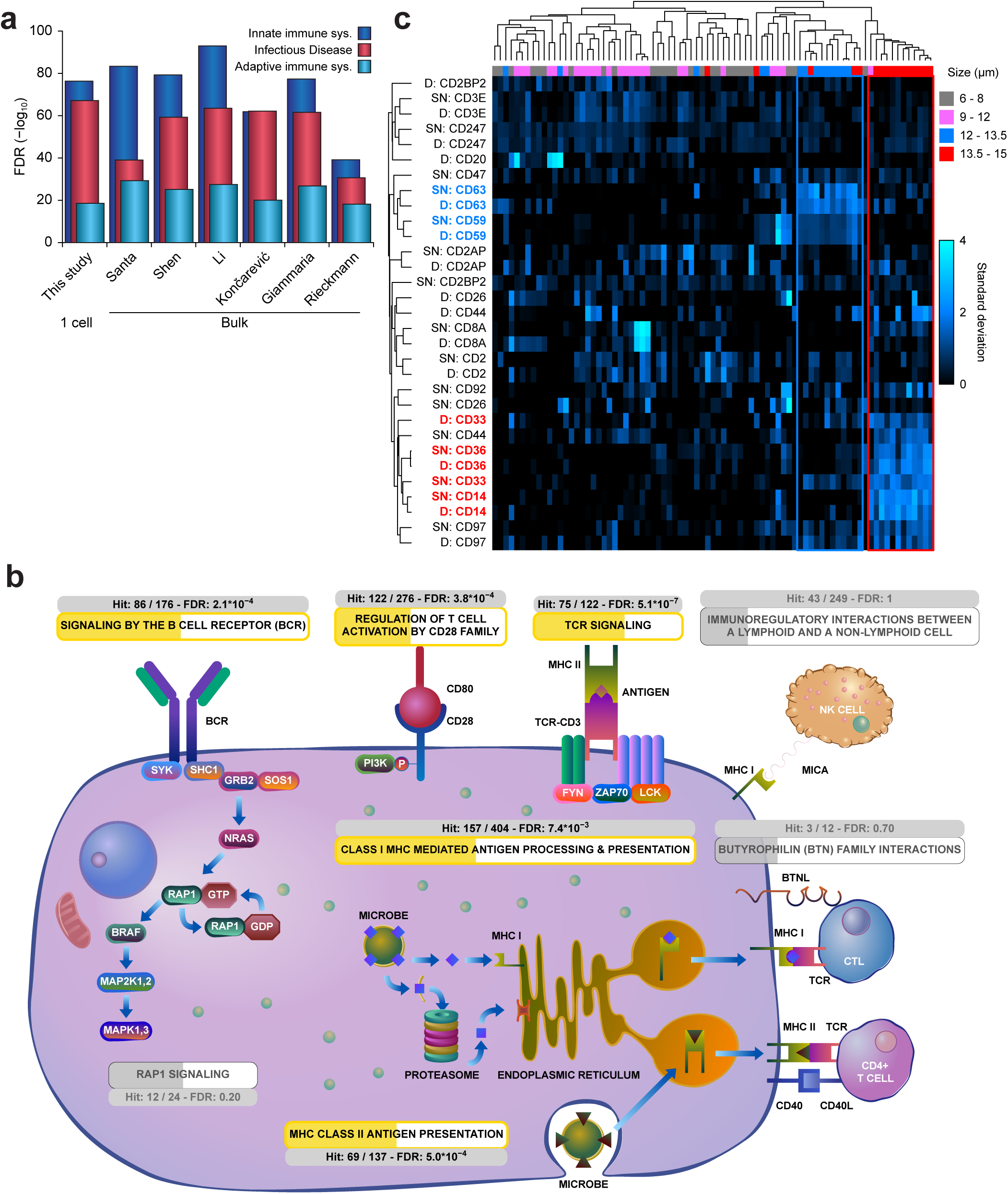
Profiling single peripheral blood mononuclear cells with proteome depth comparable to bulk analysis. Corresponds to Figure 8. (A) Reactome term enrichment analysis for the indicated immune-related Reactome Pathway categories via the STRING database (Szklarczyk et al., 2023), using all protein-coding genes identified from single PBMCs in this study (see Figure 7), or proteins as reported in six bulk PBMC studies (Giammaria et al., 2025; Koncarevic et al., 2014; Li et al., 2022; Rieckmann et al., 2017; Santa et al., 2024; Shen et al., 2019). The inverse log_10_ false discovery rate (FDR) corresponding to each category is plotted, with higher values indicating a higher degree of statistical significance. (B) Reactome Pathway enrichment analysis performed using and curated by Reactome (Milacic et al., 2024), using all protein-coding genes identified from single PBMCs as input, and visualizing the Adaptive Immune System pathway. The numbers of hits within each sub-pathway are indicated, along with FDR as calculated by Reactome. Significantly over-represented pathways within our single PBMC analysis are highlighted in gold. (C) Unsupervised hierarchical Pearson clustering of all CD marker proteins identified in the Mixed PBMC population (Figure 7D). The two most distinct cellular clusters of CD markers are highlighted, boxed and colored in red and blue. Sizes of all individual cells, as recorded by the CellenOne during sorting, are indicated. Cell size was not used to drive clustering. *n* = 81 single PBMCs. “D”; identification and quantification via DIA-NN, “SN”; identification and quantification via Spectronaut.

## SUPPLEMENTARY NOTE LEGENDS

**Supplementary Note 1.**
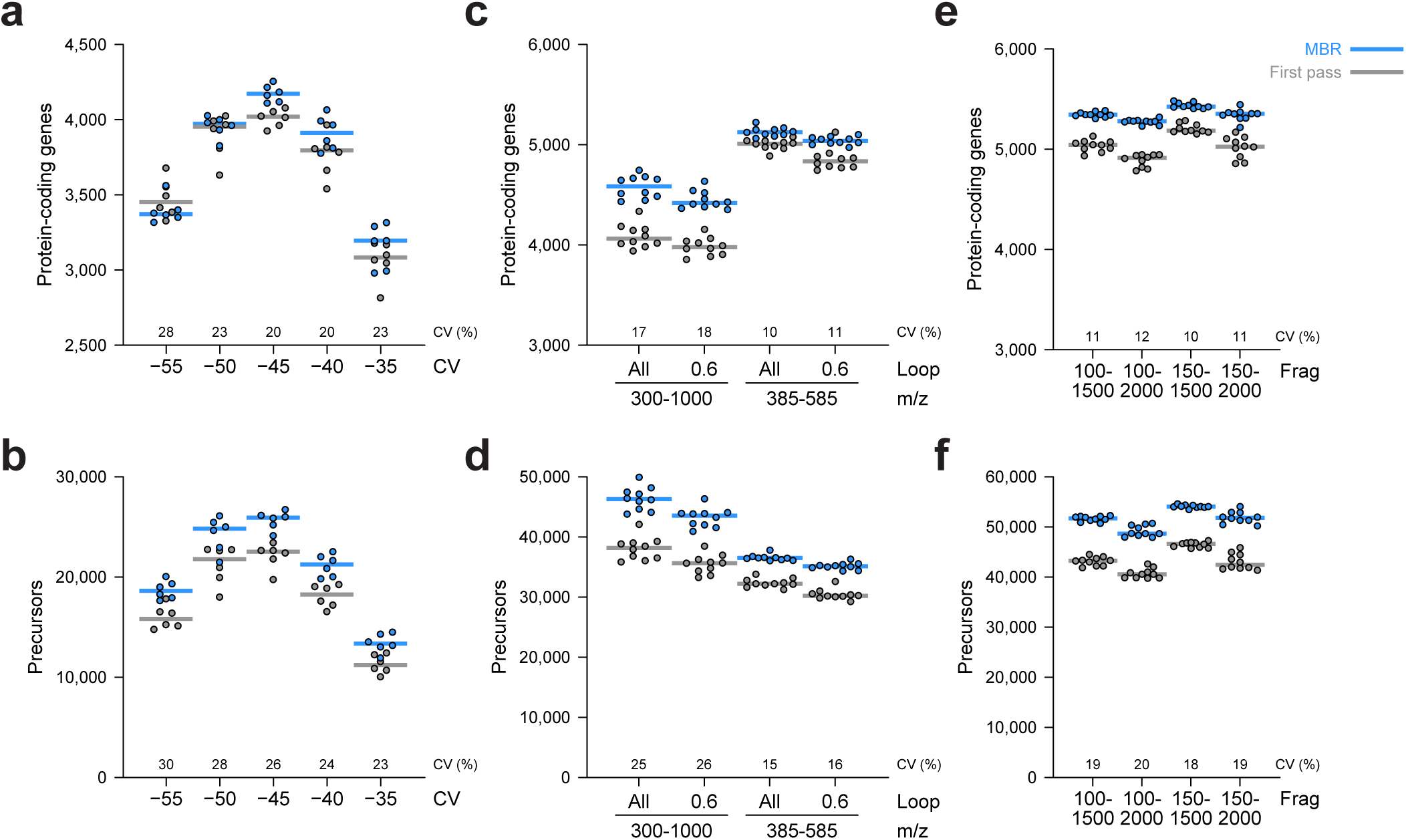
Further optimization of MS-technical method parameters. (A) Number of protein-coding genes identified when analyzing samples with a high-field asymmetric-waveform ion-mobility spectrometry (FAIMS) device, at the indicated compensation voltages (CV; on x-axis), for analysis of 250 pg HeLa. All instrument settings are detailed in Supplementary Table X. Bars represent median, with gray and blue corresponding to data analysis using DIA-NN in first pass mode and MBR mode, respectively. The numbers above the x-axis represent median coefficient of variation percentage, *n* = 6 technical replicates. (B) As **A**, but showing number of precursors. (C) As **A**, but comparing the “All” and 0.6 seconds loop control settings, for both the 300-1,000 *m*/*z* and 385-585 *m*/*z* scanning methods, for analysis of 250 pg HeLa without FAIMS installed. *n* = 10 technical replicates. (D) As **C**, but showing number of precursors. (E) As **A**, but comparing different fragment recording ranges (“Frag”), for analysis of 250 pg HeLa without FAIMS installed. *n* = 10 technical replicates. (F) As **E**, but showing number of precursors.

**Supplementary Note 2.**
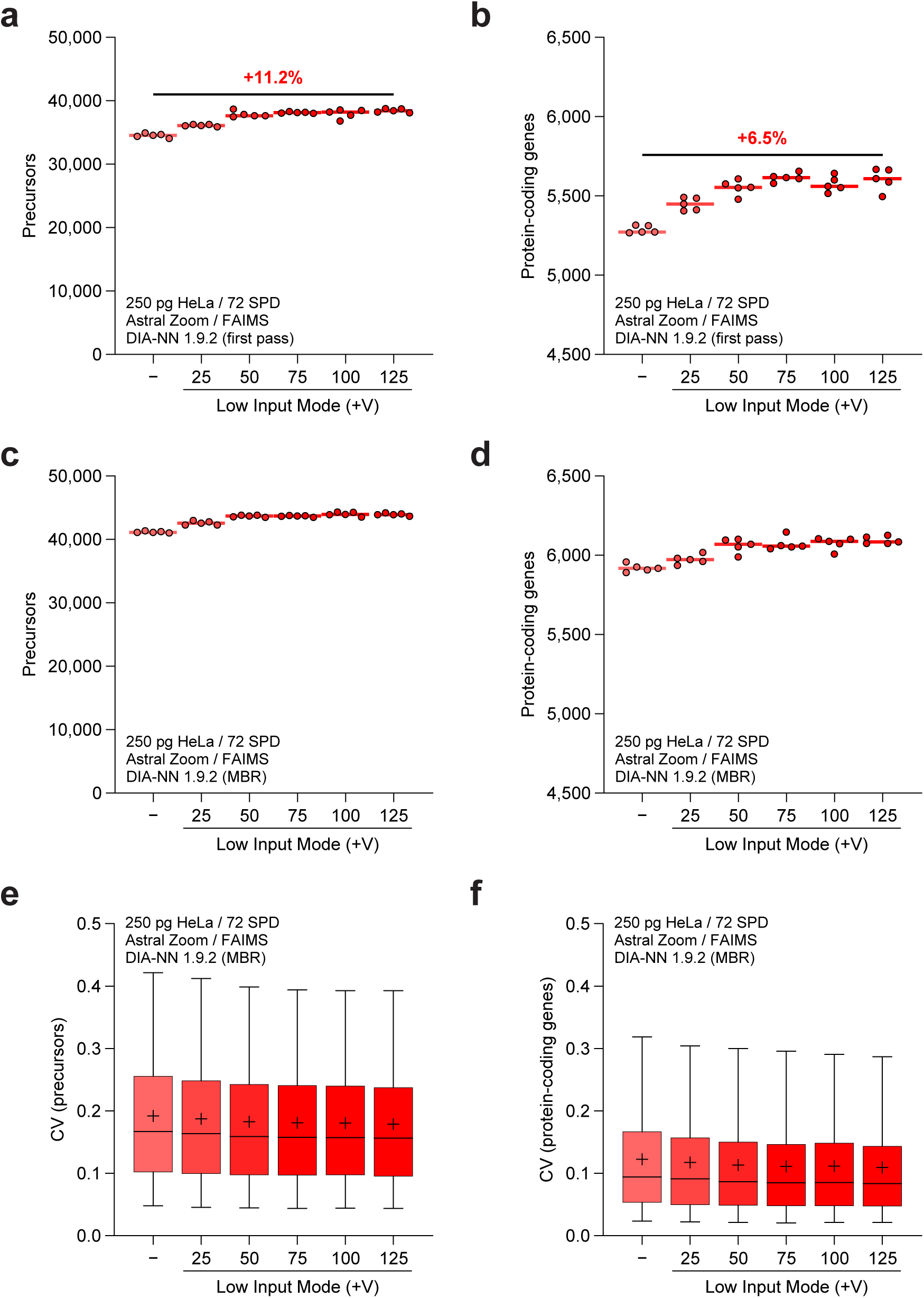
Evaluation of Low Input mode on the Orbitrap Astral Zoom MS. (A) Number of precursors identified from 250 pg HeLa lysate when utilizing standard analysis mode or a manually adjusted PMT voltage to emulate Low Input mode. Bars represent median, percentage increase of median value is indicated. Fata analysis using DIA-NN v1.9.2 in first pass mode, *n* = 5 technical replicates. (B) As **A**, but for protein-coding genes. (C) As **A**, but using matching-between-runs (MBR) with DIA-NN. (D) As **B**, but using MBR. (E) Coefficient of variation for all identified precursors as a box plot. Central line; median, plus symbol; average, box limits; 1^st^ and 3^rd^ quantile, whisker limits; 5^th^ and 95^th^ percentile. Data analysis using DIA-NN v1.9.2 with matching-between-runs, with quantification at the MS/MS level. (F) As **E**, but for protein-coding genes.

**Supplementary Note 3.**
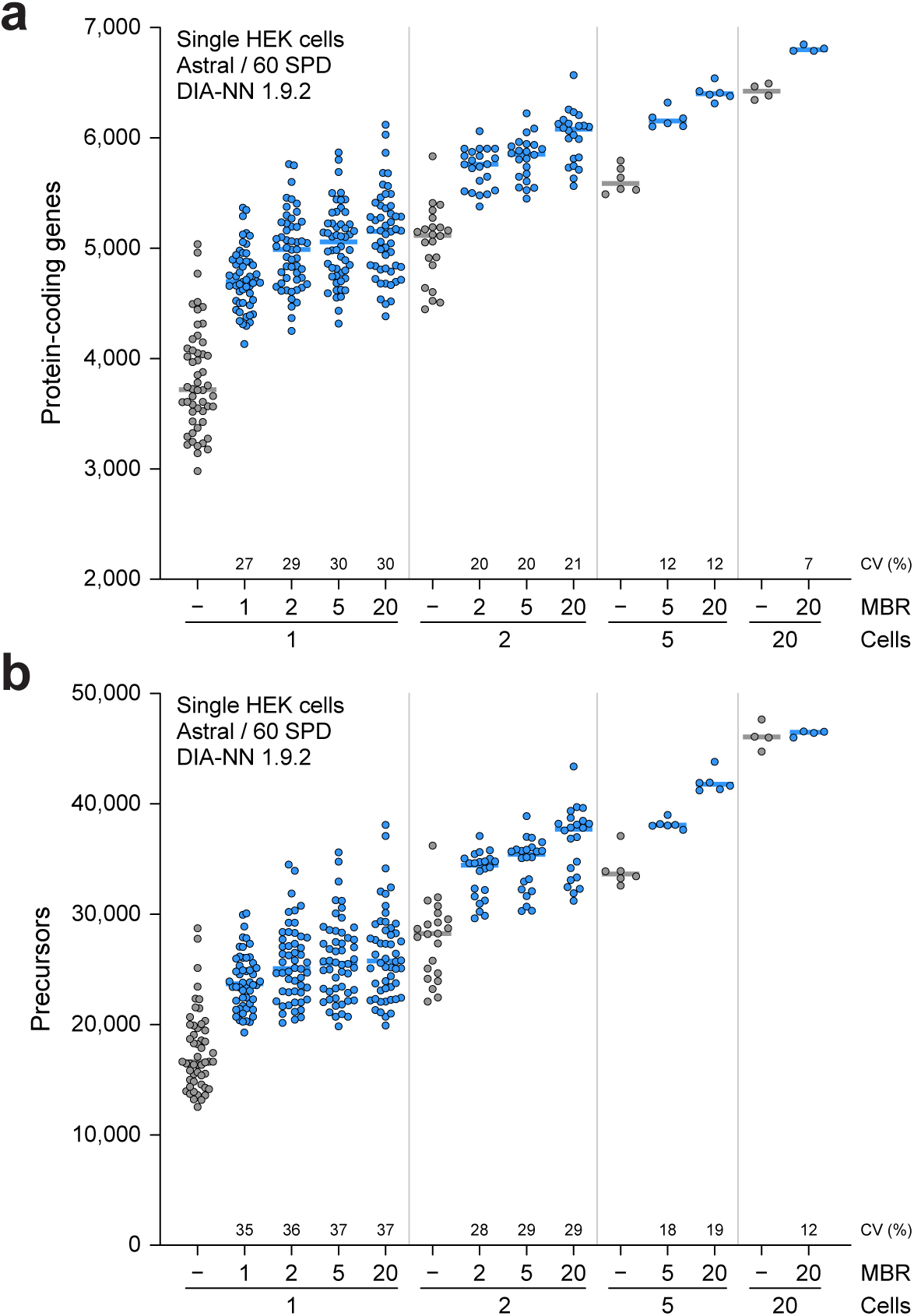
Evaluation of the carrier effect when analyzing single or multiple HEK cells. (A) Overview of the number of protein-coding genes identified from single or multiple HEK cells, following fully automated handling and digestion using the CellenOne (see also Figure 4A). Cells were sorted as 1 cell, 2 cells, 5 cells, or 20 cells, and analyzed using an Orbitrap Astral MS. Data analysis was performed using DIA-NN, either in first pass mode (in gray) or using matching-between runs (in blue) to all other 1 cell, 2 cell, 5 cell, or 20 cell runs, as indicated. Bars represent median. Numbers above the x-axis are median coefficient of variation. *n* = 52 (×1 cell), 22 (×2 cells), 6 (×5 cells), 4 (×20 cells). (B) As **A**, but for precursors.

